# RNA-Seq reveals that *Pseudomonas aeruginosa* mounts growth medium-dependent competitive responses when sensing diffusible cues from *Burkholderia cenocepacia*

**DOI:** 10.1101/2023.02.11.528112

**Authors:** Anne Leinweber, Clémentine Laffont, Martina Lardi, Leo Eberl, Gabriella Pessi, Rolf Kümmerli

## Abstract

Most habitats host diverse bacterial communities, offering opportunities for inter-species interactions. While competition might often dominate such interactions, little is known about whether bacteria can sense competitors and mount adequate responses. The competition-sensing hypothesis proposes that bacteria can use cues such as nutrient stress and cell damage to prepare for battle. Here, we tested this hypothesis by measuring transcriptome changes in *Pseudomonas aeruginosa* exposed to the supernatant of its competitor *Burkholderia cenocepacia*. We found that *P. aeruginosa* exhibited significant and growth-medium-dependent transcriptome changes in response to competition. In iron-rich medium, *P. aeruginosa* up-regulated genes encoding the type-VI secretion system and the siderophore pyoverdine, whereas genes encoding phenazine toxins and hydrogen cyanide were upregulated under iron-limited conditions. Moreover, general stress response and quorum-sensing regulators were upregulated upon supernatant exposure. Altogether, our results reveal nuanced competitive responses of *P. aeruginosa* when confronted with *B. cenocepacia* supernatant, integrating both environmental and social cues.

## Introduction

Sequencing technologies have revealed that natural bacterial communities are incredibly diverse, and that myriads of taxa can live in close proximity to one another ^1–3^. From an ecological perspective, the living together of different species can lead to both cooperative and competitive interactions. While cooperative interactions often involve cross-feeding of primary and secondary metabolites ^4–6^, intense inter-specific competition for essential resources such as nutrients, space and oxygen prevail in many habitats ^7–11^. This view is supported by the fact that many bacterial species have evolved mechanisms to attack and defend themselves from competitors ^12,13^. Defensive strategies include the formation of protective biofilms, motility to move away from competitors or the use of efflux pumps to eject toxic compounds ^14–19^. Attacking measures typically involve toxins that are either secreted in the environment, or directly injected into competitor cells via contact-dependent secretion systems ^13,20–28^.

Recently, there has been much interest in understanding whether and how bacteria can sense the presence of competitors and mount adequate responses ^29–36^. This interest roots in the evolutionary thinking that defensive and attacking responses are costly and should only be mounted when required. Cornforth and Foster ^29^ developed a concept for competition sensing by proposing that bacteria rely on cues such as resource limitation or competitor-induced cell damage, to trigger competitive counter measures such as toxin production and biofilm formation. Moreover, it was proposed that bacteria might be able to sense competitors via their secreted diffusible molecules, such as toxins or quorum-sensing molecules, to mount responses before they physically meet the competitor ^30,32^.

Here we used transcriptomics to test whether *P. aeruginosa* PAO1 ^37^ changes its gene expression profile when exposed to the supernatant of a competitive species, *B. cenocepacia* H111, containing secreted compounds that could serve as cues for competition sensing. We chose these two bacteria as a model system because both species are opportunistic human pathogens, which can co-infect lungs of cystic fibrosis patients, but potentially also co-exist in natural soil habitats ^38–44^. We thus reasoned that *P. aeruginosa* might have evolved ways to sense and respond to diffusible cues from *B. cenocepacia*. To test our hypothesis, we exposed *P. aeruginosa* to spent supernatant from *B. cenocepacia* cultures mixed with fresh medium and compared its transcriptome to conditions where it grew in fresh medium alone or when exposed to its own supernatant (Fig. 1). We carried out this setup in three-fold replication under both iron-rich and iron-limited conditions. We then used the replicated transcriptomic data (n = 18) to perform the following main analyses. First, we compared the transcriptomes of *P. aeruginosa* growing in iron-rich versus iron-limited casamino acid (CAA) medium without supernatant supplementations to validate whether genes known to be iron-regulated respond to our treatment in the expected way. Second, we conducted a global analysis to quantify the number and type of differentially up-and downregulated genes across the supernatant treatments in the two nutrient environments. Third, we focused on specific gene classes encoding competitive traits and regulators that could potentially be involved in competition sensing and tested whether they are differentially expressed when *P. aeruginosa* is exposed to the supernatant of *B. cenocepacia*.

**Fig 1.**
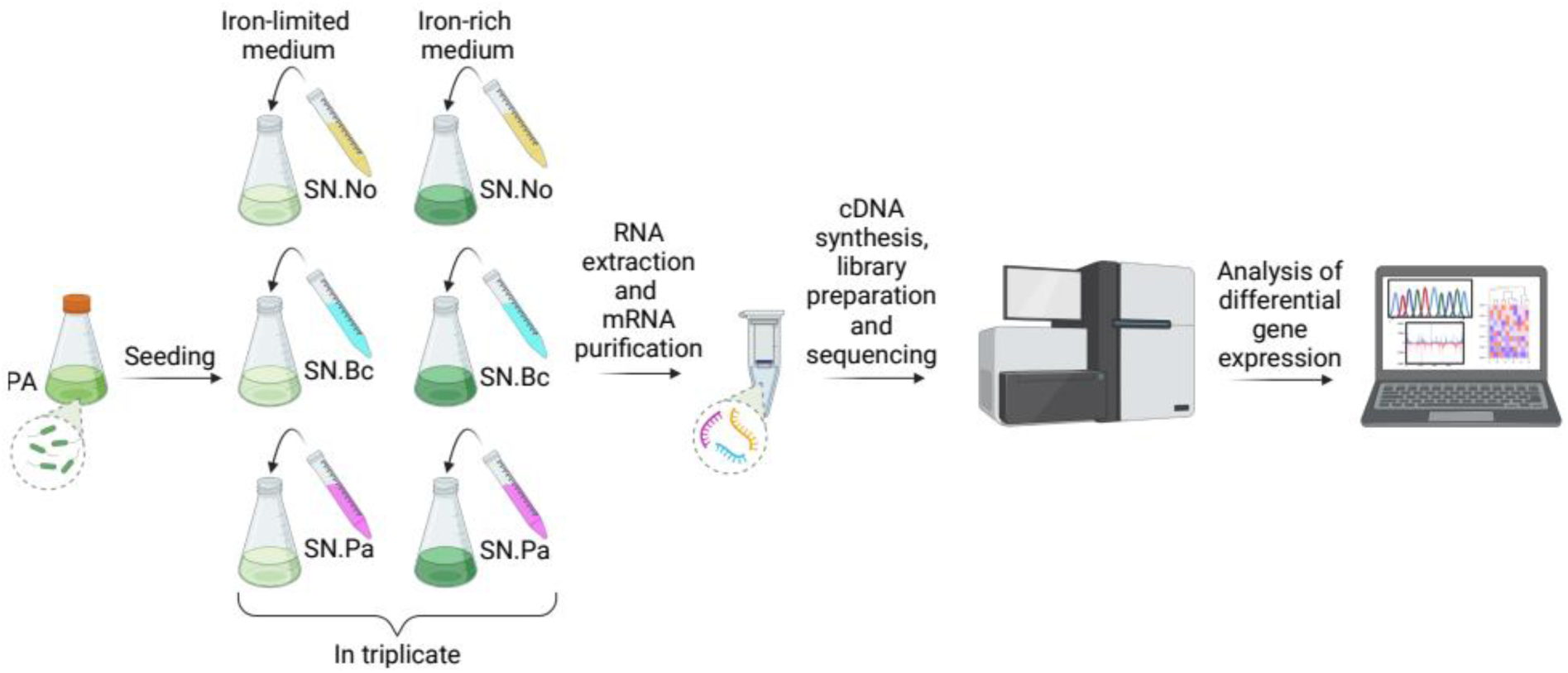
Experimental design and workflow of our transcriptomic study. Starter cultures of *P. aeruginosa* PAO1 were grown under standardized conditions, harvested upon reaching the late exponential phase, and prepared to seed the experimental treatments in triplicates. We had three main treatments under both iron-limited and iron-rich conditions. Yellow (SN.No): *P. aeruginosa* grows in 100% fresh casamino acid (CAA) medium. Cyan (SN.Bc): *P. aeruginosa* grows in 70% fresh CAA + 30% of spent supernatant of its competitor *B. cenocepacia* H111. Magenta (SN.Pa): *P. aeruginosa* grows in 70% fresh CAA + 30% of its own spent supernatant. Upon reaching the last third of the exponential growth phase, cells were pelleted and their RNA extracted and processed for transcriptomics analysis (RNA preparation and cDNA synthesis, library preparation, sequencing and gene expression analyses). Culture growth was closely monitored (Supplementary Fig. 1) to ensure that cultures were harvested before they experienced severe nutrient limitation. This figure was created with BioRender.com.

## Results

### *P. aeruginosa* transcriptomes segregate in response to supernatant treatment and iron availability

We grew *P. aeruginosa* in 100% fresh CAA medium (no supernatant = SN.No) and in 70% fresh CAA medium supplemented with either 30% of its own spent supernatant (*P. aeruginosa* supernatant = SN.Pa) or 30% of the spent supernatant of its competitor (*B. cenocepacia* supernatant = SN.Bc). The three growth treatments were carried out under both iron-rich and iron-limited conditions (Fig. 1). Across the six conditions and the three replicates per condition, we could map between 4 595 376 and 8 827 668 reads to the PAO1 genome (Supplementary Table 1). Based on these reads, we conducted a principal component analysis (PCA) using the rlog transformed and normalized raw counts data ^45^. We found that the first three principal components (PC) explained 75.2 % of the total variance observed (Fig. 2a+b). While PC1 reveals a clear separation between the cultures growing under iron-rich versus iron-limited conditions, PC3 shows segregation between cultures growing in 100% CAA medium and cultures that received spent supernatant in addition to 70% CAA. We further observed that the triplicates of the same treatment clustered together, demonstrating that our independent biological replicates yielded reproducible results. The strong clustering of replicates and the distinct differences in the transcriptomes between iron-rich and iron-limited conditions also emerged in a Euclidean distance matrix analysis (Fig. 2c).

**Fig. 2.**
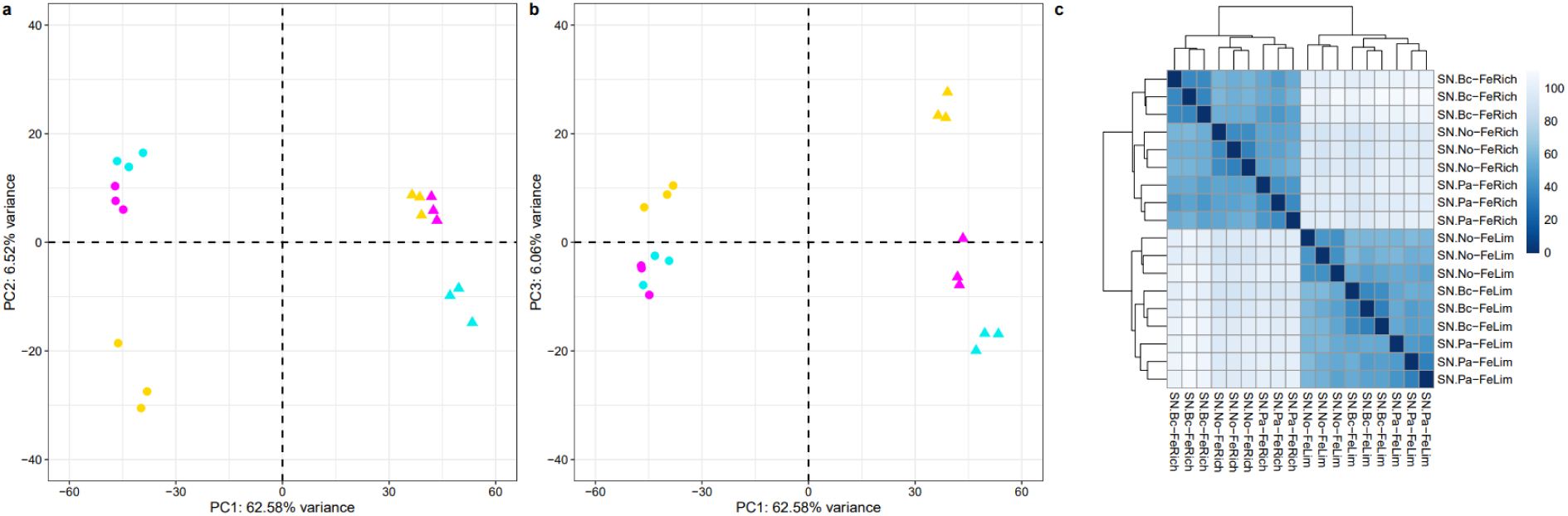
Principal component and Euclidean distance analyses reveal separation between experimental conditions and clustering between biological replicates of the same condition. (**a**) + (**b**) The first three principal components (PC) on all transcriptomics read explain 75.2 % of all the variation observed in the data set. Circles and triangles depict replicates grown under iron-limited and iron-rich conditions, respectively. Yellow symbols: *P. aeruginosa* grown in 100% fresh casamino acid (CAA) medium (SN.No). Cyan symbols: *P. aeruginosa* grown in 70% fresh CAA + 30% of spent supernatant of its competitor *B. cenocepacia* H111 (SN.Bc). Magenta symbols: *P. aeruginosa* grown in 70% fresh CAA + 30% of its own spent supernatant (SN.Pa). (**c**) Pairwise Euclidean distance analysis between all replicates, shown as a heatmap covering the scale from 0 (absolute identity, dark blue) to the maximum possible discrepancy (white). PC and Euclidean distance analyses both reveal a clear separation between replicates grown in iron-limited (FeLim) versus iron-rich (FeRich) conditions and a clustering of the three replicates grown under the same supernatant condition.

### Expression of siderophore genes and other iron-regulated genes respond to iron limitation

To validate our transcriptomic data, we tested whether the expression of genes known to be regulated by iron availability responded in the expected way in our conditions where bacteria grew in plain CAA medium (iron-limited versus iron-rich) without the addition of supernatants (Fig. 3). We first focused on the genes involved in the synthesis, secretion and regulation of the two iron-scavenging siderophores pyoverdine and pyochelin ^46–48^. As expected, we observed that all the 30 siderophore genes (Fig. 3) were significantly upregulated under iron limitation. The expression of *toxA* and *prpL* (*piv*), encoding exotoxinA and protease IV, which are known to be induced in the presence of pyoverdine ^49,50^, were also significantly upregulated under iron limitation. Next, we focused on two genes encoding the superoxide dismutase SodB (iron-dependent) and SodM (manganese-dependent).

**Fig. 3.**
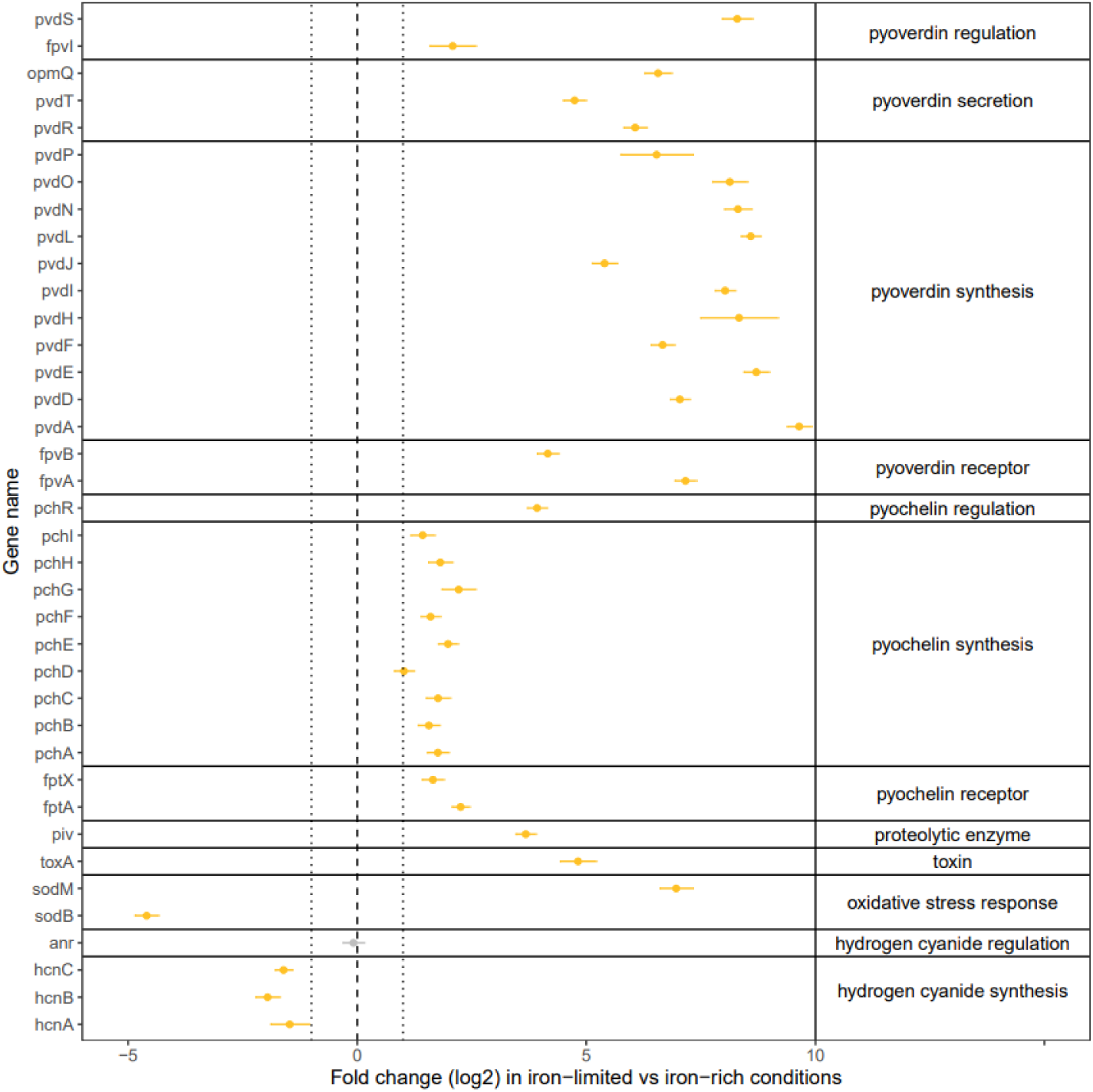
Gene expression analysis reveals that genes known to be iron regulated respond to iron limitation. Log2-fold change (iron-limited versus iron-rich CAA medium) in the expression of a set of representative genes known to be regulated in response to iron availability. The data shown are from the SN.No conditions where *P. aeruginosa* was grown in 100% CAA medium without supernatants, supplemented with either no iron or 20 μM FeCl_3_. Data represent the means and standard deviations from the three biological replicates. Yellow and grey dots depict genes that are significantly and not-significantly differently regulated, respectively (FDR-p_adj_ < 0.05). Genes are grouped according to the functional unit they belong to.

We observed that *sodB* is downregulated, whereas *sodM* is upregulated under iron limitation, a pattern that is in line with the transcriptional switch described by Bollinger et al. ^51^. Finally, we examined the expression of the genes involved in hydrogen cyanide synthesis, which is controlled by the ANR regulator featuring an iron-sulfur cluster ^52,53^. Consistent with previous work, we found that *anr* expression itself was not affected by iron availability, but the expression of *hcnABC* decreased under iron limitation. These control analyses confirm that our RNA-Seq analysis yields reliable results. Finally, we also measured the production of the siderophore pyoverdine over time as a key phenotype affected by iron availability. Consistent with our transcriptomic data, we found higher levels of pyoverdine production in iron-limited compared to iron-rich conditions (Supplementary Fig. 2). In line with previous work, we further observed that *P. aeruginosa* down-scaled *de novo* pyoverdine production when growing in its own supernatant, which already contains pyoverdine ^54^.

### *P. aeruginosa* shows multiple distinct changes in gene expression when exposed to the supernatant of *B. cenocepacia*

Next, we examined differential gene expression of *P. aeruginosa* between our three growth conditions (no supernatant = SN.No, *P. aeruginosa* supernatant = SN.Pa and *B. cenocepacia* supernatant = SN.Bc), separately for iron-rich and iron-limited conditions (Fig. 4). We considered a gene to be differentially expressed with log2-fold change values > 1 or < -1 and with FDR-adjusted p-values < 0.05. Our RNA-Seq analysis revealed that > 98% out of the 5713 known genes were expressed (i.e. we found at least one read; Supplementary Table 2). Across all pairwise comparisons, between 5.7% and 10.7% of the genes were differentially expressed, of which on average 3.3% and 4.4% were up-and downregulated, respectively (Supplementary Table 2). The Venn diagrams in Fig. 4a+b show the number of differentially expressed genes that are unique or shared across the growth conditions.

We then compared the gene expression profile of *P. aeruginosa* cells grown with *B. cenocepacia* supernatant (SN.Bc) to the no (SN.No) and own (SN.Pa) supernatant condition. We had four comparisons in total (SN.Bc vs. SN.No and SN.Bc vs. SN.Pa in both iron-limited and iron-rich medium), and we show the MA plots (Fig. 4c-f), a functional classification of the differentially regulated genes (Fig. 4g-j), and the top 50 up-and downregulated genes (Supplementary Table 3-6) for all four comparisons. We observed more gene COG-categories to be significantly over- or underrepresented in the SN.Bc vs. SN.No comparison than in the comparison between the two supernatant conditions (SN.Bc vs. SN.Pa). This finding agrees with our observation that the SN.Bc and SN.Pa conditions are close to one another in the PCA plot (Fig. 2) and that only a relatively small number of genes were differentially regulated in this comparison (Fig. 4 a+b). To give a complete picture and to demonstrate how *P. aeruginosa* changes gene expression in response to its own supernatant, we repeated the above analysis for the third comparison (SN.Pa vs. SN.No) and show it in Supplementary Fig. 3a-d.

When focusing on the SN.Bc vs. SN.No comparison (showing more pronounced differences), we found four defined gene COG-categories to be significantly overrepresented among the upregulated genes under iron-rich conditions (Fig. 4g): (E) amino acid transport and metabolism, (I) lipid transport and metabolism, (P) inorganic ion transport and metabolism, and (Q) secondary metabolite biosynthesis transport and catabolism. Especially the differential regulation of genes involved in secondary metabolites production is interesting as they often encode competitive traits. Indeed, the table of the top 50 differentially regulated genes (Supplementary Table 3) features genes involved in the synthesis of the secondary metabolites pyoverdine, rhamnolipid, and phenazines. Under iron-limited conditions, seven gene COG-categories were significantly overrepresented among the upregulated genes (Fig. 4i). The COG-categories predominantly involve metabolism: amino acids (E), nucleotides (F), carbohydrates (G), coenzymes (H), lipids (I); but also energy conversion (C) and transcription (K). These results suggest that *P. aeruginosa* increases metabolic activities in the presence of *B. cenocepacia* supernatant under iron-limiting conditions, either because the medium composition changed or as a direct competitive response against *B. cenocepacia*.

Beyond these global analyses, two observations in Supplementary Tables 3-6 caught the eye. First, genes of the PA0878-PA0883 cluster were among the topmost upregulated genes in iron-rich conditions (Supplementary Tables 3 and 5). These genes encode enzymes involved in the degradation of mesaconate and itaconate ^55,56^. The two acids show antimicrobial activities and are produced by macrophages but also certain bacteria through the glutamate metabolism pathway ^57,58^. The upregulation of these genes could thus represent a defensive response against an unknown antibacterial compound in the supernatant of Bc H111. Second, genes of the clusters PA5081-PA5086 and PA1343-1334 were among the topmost upregulated genes in iron-limited conditions (Supplementary Tables 4 and 6). These genes encode proteins involved in the utilization of D-glutamic acid (D-Glu) and D-glutamine (D-Gln), two essential components of the bacterial cell wall ^59^. Their upregulation could potentially be a response to cell wall damage induced by compounds secreted by *B. cenocepacia,* although this hypothesis remains speculative as other non-competitive triggers could also be involved.

**Fig. 4.**
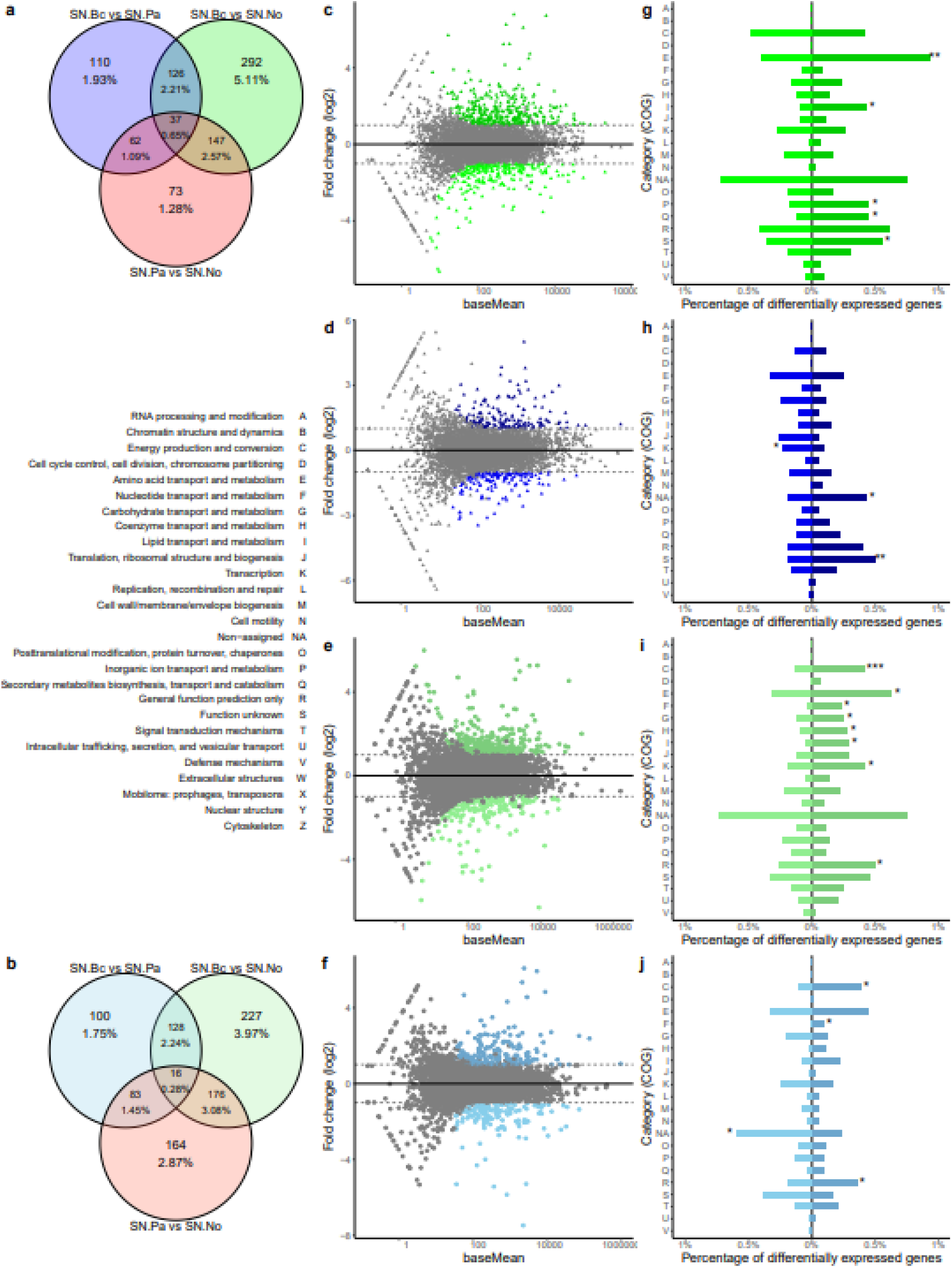
Global differential gene expression analysis of *P. aeruginosa* in response to the exposure to spent supernatant of its competitor *B. cenocepacia*. Transcriptomic analysis in triplicates were carried out for three different conditions: *P. aeruginosa* growing in (i) 100% casamino acid (CAA) medium (SN.No), (ii) 70% CAA + 30% of spent supernatant of *B. cenocepacia* H111 (SN.Bc), and (iii) 70% CAA + 30% of its own spent supernatant (SN.Pa). Differential gene expression analysis were performed for all three pairwise comparisons (green: SN.Bc vs. SN.No; blue: SN.Bc vs. SN.Pa; red: SN.Pa vs. SN.No), separately for iron-rich (intense shadings) and iron-limited (faint shadings) growth conditions. Venn diagrams show the number of genes that are differentially expressed in one, two or all the comparisons for iron-rich (**a**) and iron-limited (**b**) conditions. (**c-f**) MA-plots depicting the number of significantly differentially regulated genes as a function of basemean values (FDR-p_adj_ < 0.05 and log2-fold change > |1|) for the comparisons involving SN.Bc. (**g-j**) Percentage of differentially regulated genes according to their functional (COG) classes for the comparisons involving SN.Bc. Asterisks indicate functional classes that are significantly over-or underrepresented based on the Fisher’s Exact Test for count data comparing the number of all the expressed genes with the number of the significantly up-or downregulated genes. * p < 0.05, ** p < 0.01, *** p < 0.001.

### *P. aeruginosa* upregulates competitive traits when exposed to the supernatant of *B. cenocepacia* in a medium-specific manner

We further focused on gene clusters encoding traits that are known to be involved in inter-species interactions and tested whether they are differentially regulated upon exposure to *B. cenocepacia* supernatant. Our list of 407 genes covers a broad spectrum of putatively competitive traits, including secretion systems, efflux pumps, siderophores, toxins, motility, adhesion and biofilm components (Fig. 5 + Supplementary Table 7). In addition, we included a set of housekeeping genes (*gyrA, nadB, fabD, proC, ampC, rpsL*) as negative controls, which were (as expected) not differentially regulated (Fig. 5). Numbers and names of all genes together with their log2-fold changes and adjusted p-values are listed in Supplementary Table 8 (SN.Bc vs. SN.No) and Supplementary Table 9 (SN.Bc vs. SN.Pa). We tested whether gene clusters were significantly up-or downregulated. We argue that relevant gene clusters should be differentially regulated in both comparisons to be considered as specific responses of *P. aeruginosa* to the supernatant of *B. cenocepacia*. Conversely, gene clusters that are only upregulated in the SN.Bc vs. SN.No (*B. cenopacia* supernatant vs. no supernatant), but not in the SN.Bc vs. SN.Pa (*B. cenopacia* supernatant vs. *P. aeruginosa* supernatant) comparison more likely reflect general responses to spent media. This assumption was confirmed by analyzing differential gene expression of *P. aeruginosa* in response to its own supernatant (SN.Pa) versus no supernatant (SN.No), comparisons we show in Supplementary Fig. 3e+f and Supplementary Table 10.

In iron-rich conditions, gene clusters encoding T6SS were significantly upregulated in both comparisons (Fig. 5a+b & Supplementary Table 11). The T6SS is a speargun-type of weapon deployed against other gram negatives ^27^. *P. aeruginosa* has three T6SS clusters and a closer inspection of the data revealed that only two of the three clusters (HSI-I and HSI-II) were significantly upregulated (Supplementary Table 8+9). Furthermore, the gene cluster encoding synthesis and transportation of the siderophore pyoverdine was also significantly upregulated in both comparisons (Fig. 5a+b & Supplementary Table 11). Pyoverdine can induce iron starvation in competitors like *B. cenocepacia* that lack the receptor for pyoverdine uptake ^34^ and increased production of this siderophore can be considered a competitive response ^60^. In contrast, the gene cluster encoding efflux pumps and phenazines were significantly downregulated in both comparisons (Figure 5a+b). While efflux pumps can be used to expel toxic compounds from the bacterial cell ^19^, phenazines are broad-spectrum toxins that are also involved in redox reactions ^61^.

In iron-limited conditions, the observed gene regulatory responses of *P. aeruginosa* to *B. cenocepacia* supernatant were markedly different. Here, the gene clusters encoding phenazines and hydrogen cyanide, two groups of bacterial toxins, were significantly upregulated in both comparisons (Figure 5c+d & Supplementary Table 11). Moreover, several gene clusters were significantly downregulated in both comparisons, including the ones encoding the T1SS, synthesis and transportation of the siderophore pyochelin, the production of pyocin toxins, motility (flagella, rhamnolipids, pili) as well as biofilm structures (exopolysaccharides) (Figure 5a+b & Supplementary Table 11). Taken together, our results indicate that *P. aeruginosa* adjusts its competitive strategy in response to iron availability.

We conducted two follow-up experiments to validate our main observations that *P. aeruginosa* upregulates T6SS and pyoverdine genes under iron-rich conditions, and phenazine and hydrogen cyanide genes under iron-limited conditions (see Supplementary Information). First, we constructed four reporter strains, in which we cloned the promoter region of *icmF1* (T6SS structure component gene), *pvdA* (pyoverdine synthesis gene), *phzA1* (phenazine synthesis gene), and *hcnA* (hydrogen cyanide synthesis gene) in front of a *gfp* (green fluorescent protein) reporter gene ^62^. We subjected these reporter strains to the same treatments used in the RNA-Seq experiment and followed GFP expression over time (Supplementary Fig. 4). In agreement with the RNA-Seq results, we found *icmF1* and *pvdA* to be more highly expressed in the SN.Bc compared to SN.No treatment under iron-rich but not iron-limited conditions. For *phzA1* and *hcnA*, the gene expression signals were variable across replicates and often weak, preventing a thorough quantitative analysis. That is why we carried out quantitative PCR (qPCR) as a second validation experiment. We quantified the log2-fold changes in gene expression between SN.Bc and SN.No in iron-rich and iron-limited conditions for *tssB1* (T6SS structural component gene), *pvdA*, *phzA1*, and *hcnA*. The results from the qPCR experiment validate the RNA-Seq data for all four genes (Supplementary Fig. 5).

**Fig. 5.**
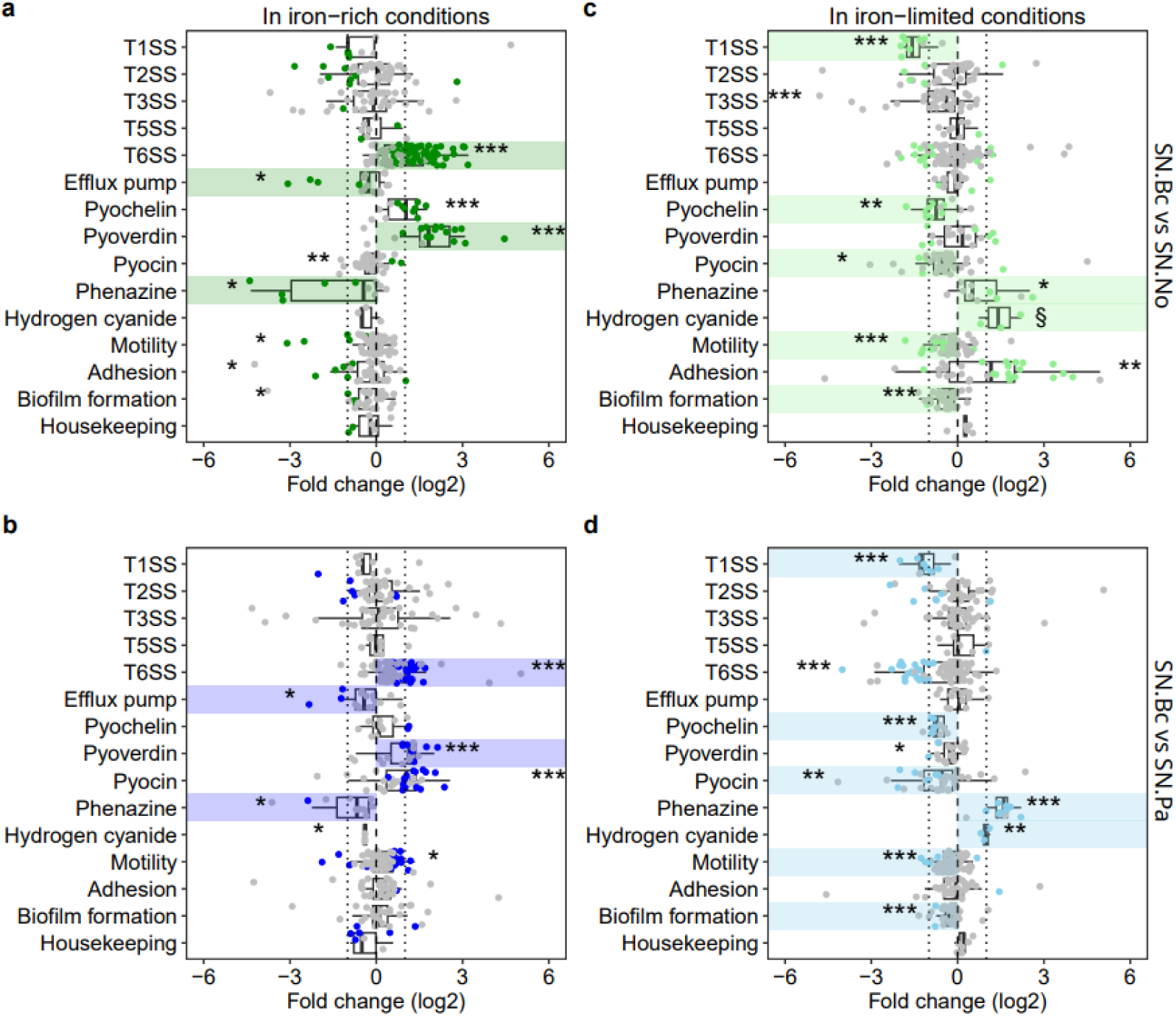
*P. aeruginosa* increases the expression of specific gene clusters encoding competitive traits when exposed to *B. cenocepacia* supernatant in an iron-dependent way. Transcriptomics analysis in triplicates were carried out for three different conditions: *P. aeruginosa* growing in (i) 100% casamino acid (CAA) medium (SN.No), (ii) 70% CAA + 30% of spent supernatant of *B. cenocepacia* H111 (SN.Bc), and (iii) 70% CAA + 30% of its own spent supernatant (SN.Pa). Data shows the log2-fold change in the gene expressions for gene clusters encoding competitive traits for two comparisons (green: SN.Bc vs. SN.No; blue: SN.Bc vs. SN.Pa), separately for iron-rich (a+b – dark shadings) and iron-limited (c+d – light shadings) growth conditions. Each dot represents an individual gene within a cluster. Colored and grey dots depict genes that are significantly and not-significantly differently regulated at the single gene level, respectively (FDR-padj < 0.05). Gene clusters highlighted in colors are those that are differentially regulated in both comparisons (SN.Bc vs. SN.No and SN.Bc vs. SN.Pa) and are considered as specific responses of *P. aeruginosa* to *B. cenocepacia*. Asterisks indicate gene clusters that are, as a group, significantly up-and downregulated, according to a t-test conducted using zero as the theoretical mean (* p < 0.05, ** p < 0.01, *** p < 0.001). § All individual genes in this group are significantly upregulated, but only marginally so when treated as a group. This is the result of the small number of genes (n=3) in this category and thus due to low statistical power. The dotted vertical lines represent log2-fold change = |1|.

### General stress response, quorum sensing, and two-component systems are involved in competition sensing

Finally, we focused on potential regulatory elements guiding competition sensing (Supplementary Tables 7-9). We followed Cornforth & Foster ^29^ who highlighted three classes of potential competition sensors: (i) general stress response components, (ii) quorum-sensing systems (Las-, Rhl-, PQS-systems), and (iii) two-component signaling systems. The three classes are not mutually exclusive. Here, we keep all the two-component signaling systems together, even though some of them (e.g. *gacS*) are involved in the general stress response ^63,64^. In Fig. 6, we plot the log2-fold changes of 93 genes belonging to these three classes in *P. aeruginosa*. Tags and names of all genes together with their log2-fold changes and adjusted p-values are listed in Supplementary Table 8 (SN.Bc vs. SN.No) and Supplementary Table 9 (SN.Bc vs. SN.Pa). Because sensors, regulators and stress response elements can act either as activators or inhibitors of specific functions, we compared the log2-fold changes of the comparison SN.Bc vs. SN.No against the comparison SN.Bc vs. SN.Pa to identify genes that were significantly up (+) or down (-) regulated in both comparisons, indicating a specific response to *B. cenocepacia* supernatant.

In iron-rich conditions (Fig. 6a), we found nine such genes, including the general stress response components *rpoS* (+) and *sodB* (-); the quorum-sensing genes *rhlR* (-), *rhlI* (-) and *lasI* (-); and the two-component signaling systems *retS* (+), *narX* (+), *dctB* (+), and *parS* (-). In iron-limited conditions (Fig. 6b), we found another set of three elements that were differentially regulated when *P. aeruginosa* was exposed to the supernatant of *B. cenocepacia*, namely *pqsA* (+), *ercS* (+), and PA3271 (+). Our analysis identifies several candidate regulatory elements that could steer competition sensing, yet the candidate elements seem to differ between iron-rich and iron-limited conditions. Moreover, we found that the regulatory responses positively correlated between the two comparisons (SN.Bc vs. SN.No and SN.Bc vs. SN.Pa) in four out of the six cases (i.e., see solid trend lines in Fig. 6), suggesting that the regulatory responses to competition are consistent, especially under iron-rich conditions.

**Fig. 6.**
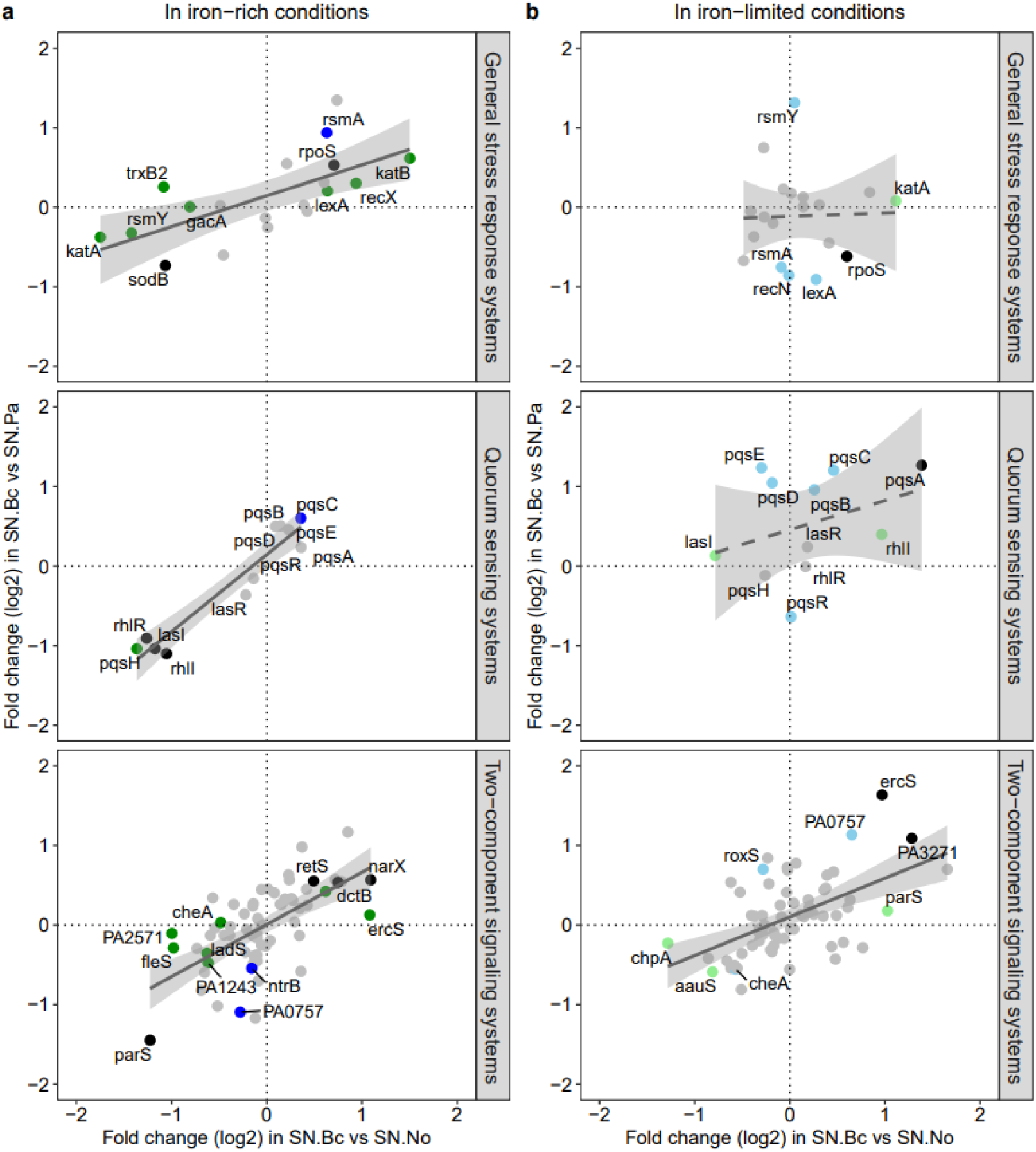
Differential expression of regulatory systems and their potential involvement in competition sensing. Panels show, from top to bottom, the log2-fold changes of genes involved in the general stress response, quorum-sensing, and two-component signaling systems. Log2-fold gene expression changes in the comparison SN.Bc vs. SN.No (*P. aeruginosa* growing in 30% of supernatant of *B. cenocepacia* vs. fresh medium without supernatant) are plotted against the comparison SN.Bc vs. SN.Pa (*P. aeruginosa* growing in 30% of supernatant of *B. cenocepacia* vs. 30% of own supernatant) for iron-rich (**a**) and iron-limited (**b**) growth conditions. Each dot depicts a regulatory gene of the corresponding class. Black dots = genes significantly differently regulated in both comparisons; grey dots = genes not significantly differentially regulated in neither of the two comparisons; green dots = genes significantly differentially regulated in the SN.Bc vs. SN.No comparison; blue dots = genes not significantly differentially regulated in the SN.Bc vs. SN.Pa comparison. Significance is based on FDR-p_adj_ < 0.05. The dashed lines, solid lines and shaded areas represent non-significant regression lines, significant regression lines (p < 0.05) and 95% confidence intervals respectively.

## Discussion

Bacteria commonly live in diverse communities, be it in environmental habitats, in infections or in host-associated microbiomes ^1,2,65^. It has increasingly been recognized that bacteria interact in these communities, with interactions covering the entire range of the social spectrum, from cooperation to commensalism to competition ^7,8^. While competition for space and nutrients seems ubiquitous, the question is whether bacteria preemptively deploy competitive traits, or whether they have evolved mechanisms to sense competitors and to mount responses when needed. The latter scenario is captured by the competition-sensing hypothesis ^29^, proposing that bacteria can use cues such as nutrient stress and competitor-induced cell damage to launch competitive responses. Here we tested this hypothesis and asked whether competition sensing is detectable at the transcriptional level in *P. aeruginosa* exposed to either its own spent medium or the supernatant of its competitor *B. cenocepacia* both in iron-rich and iron-limited medium. Our results reveal large-scale changes in the transcriptome of *P. aeruginosa*, entailing both general responses to spent medium and specific responses to the competitor. Specifically, we found genes for several competitive traits, including the T6SS, pyoverdine, phenazines, hydrogen cyanide, and putative antibacterial (e.g. itaconate/mesaconate) degrading enzymes to be upregulated upon exposure to the competitor supernatant. We further identified several genes encoding regulatory elements (general stress response, quorum sensing and two-component signaling systems) that are increasingly expressed with competitor supernatant. Moreover, transcriptional and competitive responses varied as a function of iron availability. Altogether, our results support the competition-sensing hypothesis by revealing nuanced responses of *P. aeruginosa* to *B. cenocepacia* supernatant, integrating both environmental and social information.

Transcriptomic studies, such as ours, reveal correlations between experimental treatments and shifts in gene expression patterns. They do not necessarily allow to draw direct conclusions of how exactly competition sensing works and whether changes observed are indeed caused by competitors and reflect evolutionary adaptations. Furthermore, transcriptional analysis does not take translational regulations at the protein level into account. Nonetheless, our study revealed several notable shifts in gene expression, which we wish to discuss in the context of the competition-sensing hypothesis at three different levels: (i) responses to own spent medium versus supernatant of competitor; (ii) iron-dependent responses; and (iii) regulatory elements that could steer competition sensing.

The competition-sensing hypothesis postulates that bacteria can sense competitors in two different ways: indirectly via the sensing of nutrient limitation or directly via cell damage induced by the competitor itself or its secreted molecules. Our study design allows us to partially differentiate between the two scenarios. Specifically, the SN.Bc vs. SN.No comparison captures both putative nutrient limitation (30% spent medium) and cell damage (via diffusible molecules), whereas the SN.Bc vs. SN.Pa comparison filters out limitations for basic nutrients such as carbon and nitrogen, as both treatments entail spent media. Given this setup, we identified genes of four competitive traits that were upregulated in both comparisons, suggesting that these changes reflect direct responses to the *B. cenocepacia* supernatant. Among the upregulated genes in iron-rich conditions, there were two T6SS clusters (HSI-I and HSI-II) and genes encoding the synthesis/transportation machinery of the siderophore pyoverdine (Fig. 5a+b), whereas in iron-limited conditions genes encoding the synthesis of phenazines and hydrogen cyanide were upregulated (Fig. 5c+d). T6SS, phenazines and hydrogen cyanide are well-known broad-spectrum weapons of *P. aeruginosa* against competitors ^13^. Pyoverdine can also be a competitive agent since this strong siderophore can withhold iron from competitors such as *B. cenocepacia* ^60,66^. There were also changes in gene expression that can primarily be attributed to nutrient limitation because they were only upregulated in the SN.Bc vs. SN.No comparison. The affected genes encode the synthesis/transportation machinery of the siderophore pyochelin under iron-rich condition (Fig. 5a), and several classes of metabolic genes under iron-limited conditions (Fig. 4i). In the light of the competition-sensing hypothesis, these patterns suggest that both nutrient limitation and competitor-induced cues have likely contributed to the observed gene expression changes.

We found that the putative competition-sensing responses in *P. aeruginosa* markedly differed between iron-rich and iron-limited conditions. Under iron-rich conditions, we observed the upregulation of T6SS and pyoverdine genes (Fig. 5b), whereas phenazines and hydrogen cyanide genes were upregulated under iron-limited conditions (Fig. 5d). We propose that these differential responses are the result of the different production costs associated with the respective competitive traits. T6SS is a complex structure comprising many different proteins and is likely costly to produce ^67^. Pyoverdine is also costly to produce since it involves a complex non-ribosomal peptide synthesis machinery that needs to be built up ^68^. Conversely, phenazines and hydrogen cyanide are relatively simple molecules and only few enzymes are required for their synthesis, which means that they are comparatively cheap to make ^61,69^. Thus, our finding suggests that *P. aeruginosa* invests into costly competitive traits in nutrient (e.g. iron) rich environments, whereas the bacteria switch to cheaper competitive traits when essential nutrients, such as iron, are limited. The idea that *P. aeruginosa* follows a cost-saving strategy is further supported by our observation that genes for several competitive traits are downregulated (T1SS, pyochelin, pyocins, motility, biofilm formation) under iron-limited conditions (Fig. 5c+d). Conversely, we observed that several classes of metabolic genes were significantly upregulated under iron-limited conditions (Fig. 4i). Accelerated metabolic activities is a form of nutrient competition, and our data thus indicate that *P. aeruginosa* rather invests in nutrient competition than in costly interference competition mechanisms when iron is limited.

The competition-sensing hypothesis predicts that general stress response and regulators detecting nutrient limitation and cell damage should be involved in the process ^29,35^. We indeed found that several genes involved in the general stress response ^70,71^ and two-component signaling systems ^63,64,72^ were significantly differentially regulated in both the SN.Bc vs. SN.No and SN.Bc vs. SN.Pa comparisons [iron-rich condition: *rpoS*(+), *sodB*(-), *retS*(+), *narX*(+), *dctB*(+), *parS*(-); iron-limited condition: *ercS*(+), PA3271(+); Fig. 6]. Here, we discuss three candidates, RpoS, RetS, and ParS, where links to competition-sensing may exist. RpoS is a sigma factor that mediates the general stress response in *P. aeruginosa* and other bacteria ^70,71^. It is involved in the regulation of protective measures against numerous environmental stressors like high temperature, hyperosmolarity and oxidative stress. Our finding that this key regulator is significantly upregulated in response to competition shows that our spent supernatant treatments induced significant stress in *P. aeruginosa*. ParS-ParR is a two-component system that reacts to cell wall damage induced by molecules such as polymyxin B and colistin ^73,74^. Detecting cell wall damage was highlighted as one of the most likely triggers of a competition-sensing response ^29^, which seems to be corroborated by our finding. The ParS-ParR system works both as activator and repressor ^74^, such that the observed down-regulation of *parS* could trigger the expression of an otherwise repressed trait. However, it remains unclear whether the ParS-ParR system is a repressor to any of the upregulated traits in our study. Finally, RetS is a repressor of the GacA/S two-component regulatory system ^75^. As such, it represses biofilm formation and activates T6SS synthesis ^31^. The latter matches our finding that genes encoding the T6SS were significantly upregulated. RetS is induced in response to cell lysis by kin cells and is thus a sensor for cell death ^31^, which is (similar to ParS) in support of the idea that cell damage forms the basis of competition sensing.

In contrast to iron-rich conditions, we found few stress-related regulatory candidates that could be involved in competition sensing under iron-limited conditions (Fig. 6b). Instead, we found that several genes of the PQS quorum-sensing system were significantly upregulated, particularly in the SN.Bc vs. SN.Pa comparison. The PQS quorum-sensing system directly controls the expression of phenazines and hydrogen cyanide. The increased activity of PQS genes thus matches our observation that phenazine and hydrogen cyanide genes were overexpressed. Quorum sensing was proposed to feed into competition sensing cascades ^29,76^ and our finding now lend support to this hypothesis. How the connection between quorum- and competition-sensing works yet remains an open question.

In summary, our study adds to the growing body of evidence that bacteria can sense competitors and mount specific responses to fight and withstand them ^29,34–36^. Our findings further indicate that *P. aeruginosa* reacts to both nutrient limitation and compounds secreted by the competitor *B. cenocepacia*. Another relevant insight is that the transcriptional responses observed are not uniform but highly specific to iron (nutrient) availability. Our study opens several new questions to be addressed in future research. For example, it would be interesting to test whether the same competitive traits are upregulated when *P. aeruginosa* interacts directly with *B. cenocepacia* and not just via molecules in the supernatant. It would not be surprising to see that bacteria can show nuanced responses to the sensing of diffusible molecules as opposed to direct cell contact. Another key point is whether *P. aeruginosa* can discriminate between different competitors and adjust its competitive response accordingly. At the ecological level, one might wonder how competition sensing affects species dynamics. Is competition-sensing required to displace a competitor or simply useful to defend its niche? These questions reveal that we are only at the beginning of understanding whether competition-sensing is a key modulator of bacterial interactions and whether the transcriptional changes observed in our study have at least partly evolved for the purpose of sensing competitors.

## Methods

### Bacterial strains and media

For all our experiments, we used the standard laboratory *P. aeruginosa* strain PAO1 (ATCC 15692) and *B. cenocepacia* strain H111 (LMG 23991), an isolate from a cystic fibrosis patient ^77,78^. Overnight cultures were grown in lysogeny broth (LB). All gene-expression experiments were conducted in casamino acids (CAA) medium (per 1 liter: 5 g casamino acids; 1.18 g K_2_HPO_4_·3H_2_O; 0.25 g MgSO_4_·7H_2_O) supplemented with 20 mM NaHCO_3_ and 25 mM HEPES buffer. We induced strong iron limitation by adding 100 μg/ml of the natural iron chelator human apo-transferrin. To create iron rich medium, we omitted transferrin from the above recipe and added 20 μM FeCl_3_ instead. All chemicals for the bacterial culture media were purchased from Sigma-Aldrich, Switzerland.

### Supernatant preparation from P. aeruginosa and B. cenocepacia cultures

To generate spent supernatants, we first grew cultures of *P. aeruginosa* and *B. cenocepacia* separately in LB overnight at 37°C in an orbital shaker at 220 rpm. Overnight cultures were then centrifuged at 7500 rpm for 2 minutes at room temperature, and cell pellets washed twice with 0.8% NaCl to remove spent LB medium. We then adjusted resuspended bacteria to an optical density of 1 at 600 nm (OD_600_) in 0.8% NaCl. Subsequently, we inoculated 200 µl of the bacterial suspensions (*P. aeruginosa* or *B. cenocepacia*) in 500 ml Erlenmeyer flasks containing either iron-limited or iron-rich CAA medium (150 ml) and incubated the cultures at 37°C in an orbital shaker at 220 rpm. We had three biological replicates for each species and iron treatment. We followed the growth of the *P. aeruginosa* and *B. cenocepacia* monocultures by monitoring the OD_600_ over time in a spectrophotometer (Ultrospec 2100 pro, Amersham Biosciences). After reaching stationary phase (iron-limited medium: *P. aeruginosa* at 46 h; *B. cenocepacia* at 48 h; iron-rich medium: *P. aeruginosa* and *B. cenocepacia* at 17 h), the cultures were distributed to sterile 50 ml tubes (Sarstedt) and centrifuged at 8000 rpm for 2 minutes at room temperature. Supernatants were decanted into fresh 50 ml tubes (Sarstedt) and then sterilized using PES membrane filters with low protein adsorption (0.22 µm cutoff) and stored in new 50 ml sterile tubes at -80°C until further usage.

### Experimental design of transcriptome study

For the main transcriptome experiment, we cultured *P. aeruginosa* under six different growth conditions in three-fold replication (18 cultures in total) (Fig. 1). We had one additional replicate per growth condition that we used solely to monitor growth (Supplementary Fig. 1). This is important to ensure that experimental cultures are harvested in the mid-to late-exponential phase prior to nutrient limitation (approximately 2/3 of growth; Supplementary Fig. 1). Our main treatment consisted of *P. aeruginosa* exposed to 30% of *B. cenocepacia* supernatant mixed with 70% fresh CAA medium. 30% supernatant reflects a substantial exposure of *P. aeruginosa* to *B. cenocepacia*. As control treatments, we grew *P. aeruginosa* in either 100% fresh CAA medium, or in 70% CAA medium supplemented with 30% *P. aeruginosa* supernatant. All three treatments were carried out under iron-limited and iron-rich conditions with the supernatants being generated under the same iron treatment. We first prepared two starter cultures of *P. aeruginosa* by inoculating 20 ml LB medium with 200 µl of a *P. aeruginosa* overnight culture in 100 ml Erlenmeyer flasks. These starter cultures were incubated at 37°C in an orbital shaker at 220 rpm. Once the cultures reached the late exponential growth phase, we adjusted their OD_600_ to 1, and used one of the two starter cultures to either inoculate the 12 iron-limited or the 12 iron-rich replicates. The final medium volume was 200 ml in 1l Erlenmeyer flasks, and the initial inoculum was OD_600_ = 1x10^-2^ (corresponding approximately to 9.10^6^ cells/ml). Each replicate of the *P. aeruginosa* and *B. cenocepacia* supernatant treatment contained a supernatant from an individually grown *P. aeruginosa* or *B. cenocepacia* monoculture under the respective iron condition. For culture harvesting, we followed the protocol by Pessi et al 2007 ^79^. In brief, once the cultures reached the late exponential phase, we transferred them to ice and distributed them to 50 ml tubes (Greiner) containing 5 ml ice cold stop solution (10% phenol buffered with Tris-HCl to pH8 (Sigma-Aldrich, Switzerland) and 90% ethanol). We mixed the suspension by manual inversion of tubes followed by centrifugation at 4°C for 5 minutes at 9000 rpm. We discarded the supernatant and flash-froze the cell pellet in liquid nitrogen, which was then stored at -80°C.

### RNA isolation, library preparation and RNA Sequencing

We extracted the total RNA from *P. aeruginosa* cell pellets using a modified hot acid phenol protocol (see ^79,80^ for detailed protocols), followed by the removal of short RNAs, including the 5S ribosomal RNA (5S rRNA), with the RNAeasy MiniKit (Qiagen). We completely removed the genomic DNA (gDNA) with RQ1 RNase-Free DNase (Promega), purified the RNA with the RNAeasy MiniKit (Qiagen) and the absence of gDNA was confirmed by a 40-cycles PCR using primers for the *rpoD* gene (rpoD_fwd: 5’CTGATCCAGGAAGGCAACAT; rpoD_rev: 5’TGAGCTTGTTGATCGTCTCG). The quality of the mRNA was analyzed by measuring the rRNA integrity through capillary electrophoresis in a Bioanalyzer 2100 (Agilent) using the Agilent RNA 6000 Nano Chips. We used 150 ng of RNA for cDNA synthesis and library preparation with the Encore® Complete Prokaryotic RNA-Seq DR Multiplex System from NuGEN (NuGEN Tecan, San Carlos, USA). During the cDNA synthesis, most of the ribosomal RNA in the samples was eliminated with the use of the insert-Dependent Adaptor Cleavage (lnDAC) technology to selective priming. The resulting cDNA was fragmented into 200 bp fragments with a Covaris S220 AFA system prior to the library preparation. For library preparation, we followed the protocol described by Lardi et al. ^81^. We analyzed the concentration, quality and size distribution of the libraries with capillary electrophoresis using D1000 ScreenTapestation from Agilent (size range 100-800 bp) (Agilent Technologies, Santa Clara, USA). Single-end 70 base-pair sequencing of the libraries was performed on an Illumina HiSeq2500 machine at the Functional Genomics Center Zurich.

### Analysis of differential gene expression

Illumina-generated sequencing reads were processed and mapped to the *P. aeruginosa* PAO1 genome sequence ^82^ from the *Pseudomonas* Genome Database (www.pseudomonas.com) ^83^ using the CLC Genomics Workbench v7.0 (QIAGEN CLC bio, Aarhus, Denmark). We only considered transcripts that could be unambiguously mapped (allowing up to two mismatches per read) to the *P. aeruginosa* genome. We excluded genes with no read in all the compared conditions from further analysis. The RNA-Seq raw data files of all analyzed samples are accessible through the GEO Series accession number GSE224821.

We used the Bioconductor package DESeq2 version 1.6.3 ^45^ in the R environment version 4.1.1 ^84^ for the analysis of differential gene expression. We used a principal component analysis (PCA) and a pairwise Euclidean distance analysis between all replicates to test whether our biological replicates cluster and whether there is segregation according to treatments. For this analyses, we included all expressed genes and used the regularized logarithm (rlog) transformation (which also normalizes for library size) from the DESeq2 package to increase the similarity of variances of genes with different read counts ^45^. We used the false discovery rate (FDR) test by Benjamini and Hochberg (1995) ^85^ to adjust p-values to account for multiple comparisons, and considered genes to be differentially expressed when the adjusted p-value was < 0.05 and the log2-fold change was > 1 or < -1. For some genes no p-value could be assigned. That is the case when one of the three replicates constitutes an outlier, which is determined by the Cook’s cutoff. In addition, only genes with large enough counts (depending on the basemean and variance of the expression of the single gene across the respective three replicates) generate sufficient statistical power to yield significant fold changes. To analyze the gene expression changes of *P. aeruginosa* in response to the supernatant of its competitor *B. cenocepacia*, we compared the treatment that involves this supernatant (i.e. SN.Bc) with control treatments (i.e. vs. SN.No or SN.Pa) either in iron-rich or iron-limited conditions. Firstly, we studied global changes in gene expression. Then, all the mapped genes were assigned to global functional categories defined according to the COG database ^86,87^. By using this database, some genes have been placed in several categories while other have not been assigned at all. By comparing the number of all the expressed genes with the number of the significantly up-or downregulated genes, we highlighted the functional COG-categories in which genes are differentially expressed between our conditions of interest. Secondly, we focused on 506 specific genes encoding competitive traits (i.e., secretion systems, efflux pumps, siderophores, toxins, motility, adhesion, biofilm formation) and regulators that could be potentially involved in competition sensing. For the competitive traits, we tested whether they were globally (as a group) differentially expressed in *P. aeruginosa* upon exposure to *B. cenocepacia* supernatant. Because the regulators can be activators or inhibitors, we sought to identify regulatory genes that were significantly up-or downregulated in our comparisons of interest (SN.Bc vs. SN.No and SN.Bc vs. SN.Pa).

### GFP gene expression reporter experiments

To validate the RNA-Seq results, we looked at the expression of specific genes using fluorescent gene-expression reporter strains. These strains were constructed following the protocol from Jayakumar et al.^62^. Briefly, we chromosomally integrated a transcriptional reporter fusion (in which the promoter of the gene of interest was fused to the fluorescent gene marker GFP) into the *P. aeruginosa* PAO1 wild type strain. We focused on four different genes: *icmF1* for structural component of the T6SS, *pvdA* for pyoverdine synthesis, *phzA1* for phenazine synthesis, and *hcnA* for hydrogen cyanide synthesis. By following GFP signal over time, we were able to temporally follow the expression of these genes.

The strains were grown in 10 mL of LB at 37°C and 170 rpm (Infors HT, Multitron Standard Shaker). After overnight cultures, the cells were washed in 0.8% of NaCl and OD_600_ were adjusted to 1 using a spectrophotometer (Ultrospec 2100 pro, Amersham Biosciences). 2 µL of cells were distributed into 200 µL of media in 96-well plates. The plates were incubated at 37°C in a plate reader (Tecan, Infinite Mplex). The absorbance at 600 nm and the GFP signal (excitation: 488 nm, emission: 520 nm) were measured every 15 min for 24 h with 30 sec of orbital shaking prior to each reading. We subjected bacterial cultures to the same conditions as in the RNA-Seq experiment: 100% fresh CAA medium (SN.No) and 70% fresh CAA medium supplemented with 30% of the spent supernatant of *B. cenocepacia* (SN.Bc) both in iron-rich and iron-limited conditions. The supernatants were prepared as described in the main text. In total, we had 9 or 12 replicates originating from 3 or 4 independent repeats (3 replicates per repeat) respectively in iron-rich or iron-limited conditions. We normalized all GFP signals by growth (OD + 0.01). We added 0.1 to the OD value to prevent division by zero at early time points. For statistical analysis, we calculated the area under the curve (AUC) of the normalized GFP signal over a specific time period: between 5 and 10 hours in iron-rich conditions and between 15 and 20 hours in iron-limited conditions. These time windows correspond to the approximate time points during which we harvested the cells for RNA extraction for the RNA-Seq experiment. We conducted ANOVAs to compare AUC values between SN.No and SN.Bc, after transformation of the data to meet the normality requirement. However, statistical analysis should be interpreted with caution because it is unclear at which time points gene expression differences are expected to be largest.

### qPCR gene expression experiment

The qPCR was performed on samples from *P. aeruginosa* cells exposed to fresh CAA medium (SN.No) and to *B. cenocepacia* supernatant (SN.Bc, 30% spent supernatant + 70% CAA) in iron-rich and iron-limited conditions. RNA isolation and purification was carried out from frozen cell pellets originally prepared for the RNA-Seq transcriptome study (see method above). One biological replicate (cell pellet) was used for each conditions. First, strand cDNA synthesis was performed using the M-MLV reverse transcriptase RNase H Minus (Promega) with 5 μg of total RNA from each sample and random primers (Invitro Life Technologies). cDNA was subsequently purified with the MinElute PCR Purification Kit (Qiagen). The expression of *P.aeruginosa* genes (*tssB1*, *pvdA*, *phzA1* and *hcnA*) was measured with a Stratagene MX3000P instrument (Agilent Technologies) using the Brilliant III Ultra-Fast SYBR Green qPCR Master Mix (Agilent 600883) and 0.65 μM of individual primers in a total volume of 30 μL per reaction. The primers used to target the specific genes were the following (from 5’ to 3’): tssB1_Fwd-CATCACCTTCGAGAGCA; tssB1_Rev-TCGTCGTCTTTGGGCTT; pvdA_Fwd-AGGATCACTGCGTCGTA; pvdA_Rev-AATACCACAACACCAAC; phzA1_Fwd-TGCGGATCTTCGAGAC; phzA1_Rev-TCCTTGGCGGTAGATG; hcnA_Fwd-CAGACATGACCATCCAC; hcnA_Rev-TTGCTTTCGGTTTCCAC. All qPCR reactions were performed in nine technical replicates including three cDNA dilutions (15, 7.5 and 3.25 ng) and technical triplicates for each. A low number of replicates had to be removed from the data set due to a systematic technical issue. Important to note is that here we targeted the gene *tssB1* (instead of *icmF1*) because it is more highly expressed and thus a more sensitive marker for differential T6SS expression. Fold changes were calculated using the three cDNA concentrations and based on the method from Pfaffl ^88^ without housekeeping gene normalization.

### Statistics and Reproducibility

All statistical analyses were performed with R Studio (version 4.1.1). For the analysis of differential gene expression, we used the false discovery rate (FDR) test by Benjamini and Hochberg (1995) ^85^ to adjust p-values for multiple comparisons. To investigate the global changes in gene expression, we used the Fisher’s Exact Test for count data to compare the number of all expressed genes within a functional category (from the COG database^86,87^) with the number of the significantly up- or downregulated genes in the same category. This analysis highlights the functional COG-categories in which more genes were differentially expressed between our conditions than expected by chance.

To explore whether *P. aeruginosa* shows a specific competitive response upon exposure to *B. cenocepacia* supernatant, we compared the mean of the log2-fold change of each defined competitive trait category (mean across all the genes within this category) with the expected value of zero (no differential expression) using t-tests. The sensors, regulators and stress responses were analyzed in a different way. This is because genes in these categories can be either activators or inhibitors of specific pathways and thus be up-or down-regulated, respectively. Therefore, we looked at them individually and explored whether responses to competition positively correlated across supernatant conditions (SN.Bc vs. SN.No and SN.Bc vs. SN.Pa) using linear regressions.

## Acknowledgements

We thank Stefan Wyder for help with the data analysis, Yilei Liu for support in the laboratory, and the Functional Genomics Center Zurich (FGCZ) for support with the RNA-sequencing. This project was funded by grants from the Swiss National Science Foundation (31003A_182499 to RK and 31003A_179322 to GP) and a pilot grant from the University of Zurich Research Priority Programme (URPP) Evolution in Action: from genes to ecosystems (to AL). Figure 1 was created with BioRender.com.

## Competing interests

The authors declare no competing interests.

## Data availability

All the original data of the paper will be made available upon the acceptance of this paper. The RNA-Seq raw data files of all analyzed samples are accessible through the GEO Series accession number GSE224821.

## Author contributions

AL, GP and RK designed the study. AL and CL did the lab work. AL, CL, ML and GP analyzed the data. AL, CL, ML, GP, LE and RK interpreted the data. AL, CL and RK wrote the paper with inputs from ML, GP and LE.

## Notes

### Competing Interest Statement

The authors have declared no competing interest.

### Summary of Updates

New experiments have been added as controls (figures in supplementary information).

## Literature

1. Human Microbiome Project Consortium. Structure, function and diversity of the healthy human microbiome. Nature 486, 207–214 (2012).

2. Sunagawa, S., et al. Structure and function of the global ocean microbiome. Science 348, 1261359 (2015).

3. Taş, N., et al. Metagenomic tools in microbial ecology research. Curr. Opin. Biotechnol. 67, 184– 191 (2021).

4. Giri, S., et al. Metabolic dissimilarity determines the establishment of cross-feeding interactions in bacteria. Curr. Biol. CB 31, 5547–5557.e6 (2021).

5. Rakoff-Nahoum, S., Foster, K. R. & Comstock, L. E. The evolution of cooperation within the gut microbiota. Nature 533, 255–259 (2016).

6. Kramer, J., Özkaya, Ö. & Kümmerli, R. Bacterial siderophores in community and host interactions. Nat. Rev. Microbiol. 18, 152–163 (2020).

7. Hibbing, M. E., Fuqua, C., Parsek, M. R. & Peterson, S. B. Bacterial competition: surviving and thriving in the microbial jungle. Nat. Rev. Microbiol. 8, 15–25 (2010).

8. West, S. A., Diggle, S. P., Buckling, A., Gardner, A. & Griffin, A. S. The Social Lives of Microbes. Annu. Rev. Ecol. Evol. Syst. 38, 53–77 (2007).

9. Stubbendieck, R. M., Vargas-Bautista, C. & Straight, P. D. Bacterial Communities: Interactions to Scale. Front. Microbiol. 7, (2016).

10. Bertness, M. & Callaway, R. Positive Interactions in Communities. Trends Ecol. Evol. 9, 191–193 (1994).

11. Dunny, G. M., Brickman, T. J. & Dworkin, M. Multicellular behavior in bacteria: communication, cooperation, competition and cheating. BioEssays 30, 296–298 (2008).

12. Stubbendieck, R. M. & Straight, P. D. Multifaceted Interfaces of Bacterial Competition. J. Bacteriol. 198, 2145–2155 (2016).

13. Granato, E. T., Meiller-Legrand, T. A. & Foster, K. R. The Evolution and Ecology of Bacterial Warfare. Curr. Biol. CB 29, R521–R537 (2019).

14. Hoffman, L. R., et al. Aminoglycoside antibiotics induce bacterial biofilm formation. Nature 436, 1171–1175 (2005).

15. Kaiser, D. Bacterial swarming: a re-examination of cell-movement patterns. Curr. Biol. CB 17, R561–570 (2007).

16. Marr, A. K., Overhage, J., Bains, M. & Hancock, R. E. W. The Lon protease of *Pseudomonas aeruginosa* is induced by aminoglycosides and is involved in biofilm formation and motility. Microbiol. Read. Engl. 153, 474–482 (2007).

17. Graff, J. R., Forschner-Dancause, S. R., Menden-Deuer, S., Long, R. A. & Rowley, D. C. *Vibrio cholerae* Exploits Sub-Lethal Concentrations of a Competitor-Produced Antibiotic to Avoid Toxic Interactions. Front. Microbiol. 4, 8 (2013).

18. Oliveira, N. M., et al. Biofilm Formation As a Response to Ecological Competition. PLoS Biol. 13, e1002191 (2015).

19. Burse, A., Weingart, H. & Ullrich, M. S. NorM, an *Erwinia amylovora* multidrug efflux pump involved in in vitro competition with other epiphytic bacteria. Appl. Environ. Microbiol. 70, 693– 703 (2004).

20. Bakkal, S., Robinson, S. M., Ordonez, C. L., Waltz, D. A. & Riley, M. A. Role of bacteriocins in mediating interactions of bacterial isolates taken from cystic fibrosis patients. Microbiol. Read. Engl. 156, 2058–2067 (2010).

21. Pierson, L. S. & Pierson, E. A. Metabolism and function of phenazines in bacteria: impacts on the behavior of bacteria in the environment and biotechnological processes. Appl. Microbiol. Biotechnol. 86, 1659–1670 (2010).

22. Bernier, S. P., et al. Cyanide Toxicity to *Burkholderia cenocepacia* Is Modulated by Polymicrobial Communities and Environmental Factors. Front. Microbiol. 7, 725 (2016).

23. Hou, Q., et al. Weaponizing volatiles to inhibit competitor biofilms from a distance. NPJ Biofilms Microbiomes 7, 2 (2021).

24. Mercy, C., Ize, B., Salcedo, S. P., de Bentzmann, S. & Bigot, S. Functional Characterization of *Pseudomonas* Contact Dependent Growth Inhibition (CDI) Systems. PloS One 11, e0147435 (2016).

25. Myers-Morales, T., Oates, A. E., Byrd, M. S. & Garcia, E. C. *Burkholderia cepacia* Complex Contact-Dependent Growth Inhibition Systems Mediate Interbacterial Competition. J. Bacteriol. 201, e00012–19 (2019).

26. Bleves, S., et al. Protein secretion systems in *Pseudomonas aeruginosa*: A wealth of pathogenic weapons. Int. J. Med. Microbiol. IJMM 300, 534–543 (2010).

27. Hernandez, R. E., Gallegos-Monterrosa, R. & Coulthurst, S. J. Type VI secretion system effector proteins: Effective weapons for bacterial competitiveness. Cell. Microbiol. 22, e13241 (2020).

28. Basler, M., Ho, B. T. & Mekalanos, J. J. Tit-for-tat: type VI secretion system counterattack during bacterial cell-cell interactions. Cell 152, 884–894 (2013).

29. Cornforth, D. M. & Foster, K. R. Competition sensing: the social side of bacterial stress responses. Nat. Rev. Microbiol. 11, 285–293 (2013).

30. LeRoux, M., Peterson, S. B. & Mougous, J. D. Bacterial danger sensing. J. Mol. Biol. 427, 3744–3753 (2015).

31. LeRoux, M., et al. Kin cell lysis is a danger signal that activates antibacterial pathways of *Pseudomonas aeruginosa*. eLife 4, (2015).

32. Westhoff, S., van Wezel, G. P. & Rozen, D. E. Distance-dependent danger responses in bacteria. Curr. Opin. Microbiol. 36, 95–101 (2017).

33. Westhoff, S., Kloosterman, A. M., van Hoesel, S. F. A., van Wezel, G. P. & Rozen, D. E. Competition Sensing Changes Antibiotic Production in *Streptomyces*. mBio 12, e02729–20 (2021).

34. Leinweber, A., Weigert, M. & Kümmerli, R. The bacterium *Pseudomonas aeruginosa* senses and gradually responds to interspecific competition for iron. Evol. Int. J. Org. Evol. (2018) doi:10.1111/evo.13491.

35. Lories, B., et al. Biofilm Bacteria Use Stress Responses to Detect and Respond to Competitors. Curr. Biol. CB 30, 1231–1244.e4 (2020).

36. Zarrella, T. M. & Khare, A. Systematic identification of molecular mediators of interspecies sensing in a community of 2 frequently coinfecting bacterial pathogens. PLoS Biol. 20, e3001679 (2022).

37. Granato, E. T., Ziegenhain, C., Marvig, R. L. & Kümmerli, R. Low spatial structure and selection against secreted virulence factors attenuates pathogenicity in *Pseudomonas aeruginosa*. ISME J. 12, 2907–2918 (2018).

38. Green, S. K., Schroth, M. N., Cho, J. J., Kominos, S. K. & Vitanza-jack, V. B. Agricultural plants and soil as a reservoir for *Pseudomonas aeruginosa*. Appl. Microbiol. 28, 987–991 (1974).

39. Crone, S., et al. The environmental occurrence of *Pseudomonas aeruginosa*. APMIS 128, 220–231 (2020).

40. LiPuma, J. J., Spilker, T., Coenye, T. & Gonzalez, C. F. An epidemic Burkholderia cepacia complex strain identified in soil. The Lancet 359, 2002–2003 (2002).

41. Khatun, A., et al. *Pseudomonas* and *Burkholderia* inhibit growth and asexual development of *Phytophthora capsici*. Z. Naturforschung C J. Biosci. 73, 123–135 (2018).

42. Coenye, T. & Vandamme, P. Diversity and significance of *Burkholderia* species occupying diverse ecological niches. Environ. Microbiol. 5, 719–729 (2003).

43. Aujoulat, F., et al. From environment to man: genome evolution and adaptation of human opportunistic bacterial pathogens. Genes 3, 191–232 (2012).

44. Françoise, A. & Héry-Arnaud, G. The Microbiome in Cystic Fibrosis Pulmonary Disease. Genes 11, E536 (2020).

45. Love, M. I., Huber, W. & Anders, S. Moderated estimation of fold change and dispersion for RNA-seq data with DESeq2. Genome Biol. 15, 550 (2014).

46. Escolar, L., Pérez-Martín, J. & de Lorenzo, V. Opening the Iron Box: Transcriptional Metalloregulation by the Fur Protein. J. Bacteriol. 181, 6223–6229 (1999).

47. Cornelis, P. Iron uptake and metabolism in pseudomonads. Appl. Microbiol. Biotechnol. 86, 1637– 1645 (2010).

48. Ochsner, U. A., Wilderman, P. J., Vasil, A. I. & Vasil, M. L. GeneChip expression analysis of the iron starvation response in *Pseudomonas aeruginosa*: identification of novel pyoverdine biosynthesis genes. Mol. Microbiol. 45, 1277–1287 (2002).

49. Lamont, I. L., Beare, P. A., Ochsner, U., Vasil, A. I. & Vasil, M. L. Siderophore-mediated signaling regulates virulence factor production in *Pseudomonas aeruginosa*. Proc. Natl. Acad. Sci. 99. 7072–7077 (2002).

50. Weigert, M., et al. Manipulating virulence factor availability can have complex consequences for infections. Evol. Appl. 10, 91–101 (2017).

51. Bollinger, N., Hassett, D. J., Iglewski, B. H., Costerton, J. W. & McDermott, T. R. Gene Expression in *Pseudomonas aeruginosa*: Evidence of Iron Override Effects on Quorum Sensing and Biofilm-Specific Gene Regulation. J. Bacteriol. 183, 1990–1996 (2001).

52. Blumer, C. & Haas, D. Iron regulation of the hcnABC genes encoding hydrogen cyanide synthase depends on the anaerobic regulator ANR rather than on the global activator GacA in *Pseudomonas fluorescens* CHA0. Microbiol. Read. Engl. 146 **( Pt** **10**), 2417–2424 (2000).

53. Pessi, G. & Haas, D. Transcriptional control of the hydrogen cyanide biosynthetic genes hcnABC by the anaerobic regulator ANR and the quorum-sensing regulators LasR and RhlR in Pseudomonas aeruginosa. J. Bacteriol. 182, 6940–6949 (2000).

54. Kümmerli, R. & Brown, S. P. Molecular and regulatory properties of a public good shape the evolution of cooperation. Proc. Natl. Acad. Sci. 107, 18921–18926 (2010).

55. Kronen, M., Sasikaran, J. & Berg, I. A. Mesaconase Activity of Class I Fumarase Contributes to Mesaconate Utilization by *Burkholderia xenovorans*. Appl. Environ. Microbiol. 81, 5632–5638 (2015).

56. Sasikaran, J., Ziemski, M., Zadora, P. K., Fleig, A. & Berg, I. A. Bacterial itaconate degradation promotes pathogenicity. Nat. Chem. Biol. 10, 371–377 (2014).

57. Wachsman, J. T. The role of α-ketoglutarate and mesaconate in glutamate fermentation by *Clostridium tetanomorphum*. J. Biol. Chem. 223, 19–27 (1956).

58. He, W., et al. Mesaconate is synthesized from itaconate and exerts immunomodulatory effects in macrophages. Nat. Metab. 4, 524–533 (2022).

59. He, W., Li, G., Yang, C.-K. & Lu, C.-D. Functional characterization of the dguRABC locus for D-Glu and d-Gln utilization in *Pseudomonas aeruginosa PAO1*. Microbiol. Read. Engl. 160, 2331–2340 (2014).

60. Niehus, R., Picot, A., Oliveira, N. M., Mitri, S. & Foster, K. R. The evolution of siderophore production as a competitive trait. Evol. Int. J. Org. Evol. 71, 1443–1455 (2017).

61. Laursen, J. B. & Nielsen, J. Phenazine Natural Products: Biosynthesis, Synthetic Analogues, and Biological Activity. Chem. Rev. 104, 1663–1686 (2004).

62. Jayakumar, P., Thomas, S. A., Brown, S. P. & Kümmerli, R. Collective decision-making in *Pseudomonas aeruginosa* involves transient segregation of quorum-sensing activities across cells. Curr. Biol. 32, 5250–5261.e6 (2022).

63. Sultan, M., Arya, R. & Kim, K. K. Roles of Two-Component Systems in *Pseudomonas aeruginosa* Virulence. Int. J. Mol. Sci. 22, 12152 (2021).

64. Wang, B. X., Cady, K. C., Oyarce, G. C., Ribbeck, K. & Laub, M. T. Two-Component Signaling Systems Regulate Diverse Virulence-Associated Traits in *Pseudomonas aeruginosa*. Appl. Environ. Microbiol. 87, e03089–20 (2021).

65. Tay, W. H., Chong, K. K. L. & Kline, K. A. Polymicrobial-Host Interactions during Infection. J. Mol. Biol. 428, 3355–3371 (2016).

66. Figueiredo, A. R. T., Özkaya, Ö., Kümmerli, R. & Kramer, J. Siderophores drive invasion dynamics in bacterial communities through their dual role as public good versus public bad. Ecol. Lett. 25, 138– 150 (2022).

67. Cherrak, Y., Flaugnatti, N., Durand, E., Journet, L. & Cascales, E. Structure and Activity of the Type VI Secretion System. Microbiol. Spectr. 7, 7.4.11 (2019).

68. Ringel, M. T. & Brüser, T. The biosynthesis of pyoverdines. Microb. Cell Graz Austria 5, 424–437 (2018).

69. Blumer, C. & Haas, D. Mechanism, regulation, and ecological role of bacterial cyanide biosynthesis. Arch. Microbiol. 173, 170–177 (2000).

70. Potvin, E., Sanschagrin, F. & Levesque, R. C. Sigma factors in *Pseudomonas aeruginosa* . FEMS Microbiol. Rev. 32, 38–55 (2008).

71. Suh, S. J., et al. Effect of rpoS mutation on the stress response and expression of virulence factors in *Pseudomonas aeruginosa*. J. Bacteriol. 181, 3890–3897 (1999).

72. Francis, V. I., Stevenson, E. C. & Porter, S. L. Two-component systems required for virulence in *Pseudomonas aeruginosa*. FEMS Microbiol. Lett. 364, fnx104 (2017).

73. Fernández, L., et al. Adaptive resistance to the ‘last hope’ antibiotics polymyxin B and colistin in *Pseudomonas aeruginosa* is mediated by the novel two-component regulatory system ParR-ParS. Antimicrob. Agents Chemother. 54, 3372–3382 (2010).

74. Wang, D., Seeve, C., Pierson, L. S. & Pierson, E. A. Transcriptome profiling reveals links between ParS/ParR, MexEF-OprN, and quorum sensing in the regulation of adaptation and virulence in *Pseudomonas aeruginosa*. BMC Genomics 14, 618 (2013).

75. Valentini, M. & Filloux, A. Biofilms and Cyclic di-GMP (c-di-GMP) Signaling: Lessons from *Pseudomonas aeruginosa* and Other Bacteria. J. Biol. Chem. 291, 12547–12555 (2016).

76. Niehus, R., Oliveira, N. M., Li, A., Fletcher, A. G. & Foster, K. R. The evolution of strategy in bacterial warfare via the regulation of bacteriocins and antibiotics. eLife 10, e69756 (2021).

77. Gotschlich, A., et al. Synthesis of multiple N-acylhomoserine lactones is wide-spread among the members of the *Burkholderia cepacia* complex. Syst. Appl. Microbiol. 24, 1–14 (2001).

78. Römling, U., et al. Epidemiology Of Chronic *Pseudomonas aeruginosa* Infections In Cystic Fibrosis. J. Infect. Dis. 170, 1616–1621 (1994).

79. Pessi, G., et al. Genome-wide transcript analysis of *Bradyrhizobium japonicum* bacteroids in soybean root nodules. Mol. Plant-Microbe Interact. MPMI 20, 1353–1363 (2007).

80. Liu, Y., Lardi, M., Pedrioli, A., Eberl, L. & Pessi, G. NtrC-dependent control of exopolysaccharide synthesis and motility in *Burkholderia cenocepacia* H111. PloS One 12, e0180362 (2017).

81. Lardi, M., Liu, Y., Purtschert, G., Bolzan de Campos, S. & Pessi, G. Transcriptome Analysis of *Paraburkholderia phymatum* under Nitrogen Starvation and during Symbiosis with *Phaseolus Vulgaris*. Genes 8, 389 (2017).

82. Stover, C. K., et al. Complete genome sequence of *Pseudomonas aeruginosa* PAO1, an opportunistic pathogen. Nature 406, 959–964 (2000).

83. Winsor, G. L., et al. Enhanced annotations and features for comparing thousands of *Pseudomonas* genomes in the Pseudomonas genome database. Nucleic Acids Res. 44, D646–653 (2016).

84. R Core Team. R: A language and environment for statistical computing. R Foundation for Statistical Computing, Vienna, Austria. R Found. Stat. Comput. Vienna Austria (2020).

85. Benjamini, Y. & Hochberg, Y. Controlling The False Discovery Rate - A Practical And Powerful Approach To Multiple Testing. J R. Stat. Soc Ser. B 57, 289–300 (1995).

86. Tatusov, R. L., Koonin, E. V. & Lipman, D. J. A genomic perspective on protein families. Science 278, 631–637 (1997).

87. Galperin, M. Y., Makarova, K. S., Wolf, Y. I. & Koonin, E. V. Expanded microbial genome coverage and improved protein family annotation in the COG database. Nucleic Acids Res. 43, D261–269 (2015).

88. Pfaffl, M. W. A new mathematical model for relative quantification in real-time RT–PCR. Nucleic Acids Res. 29, e45 (2001).

