## Supplementary material for "RNA-Seq reveals that *Pseudomonas aeruginosa* mounts growth medium-dependent competitive responses when sensing diffusible cues from *Burkholderia cenocepacia*": Leinweber_etal_SupplementaryInformation-BioRxive.pdf

This file contains the following supplementary information:

- 5 supplementary figures
- 11 supplementary tables

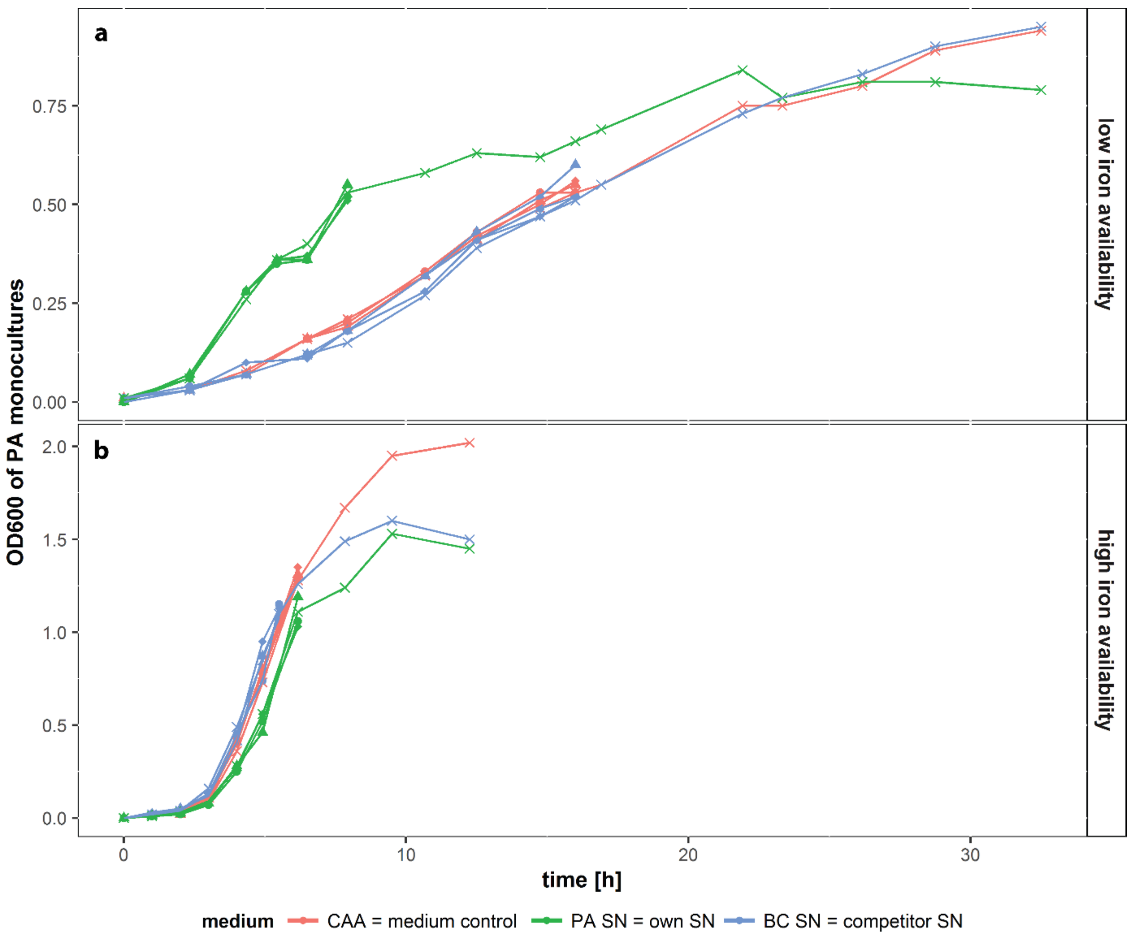


**Supplementary Fig. 1**: Monitoring growth of *P. aeruginosa* (PA) to find the right time point to harvest for following RNA isolation. PA grew in 100% fresh CAA medium (medium control, red), in 70% fresh CAA medium mixed with 30% of its own (PA) supernatant (green) or mixed with 30% of competitor (*B. cenocepacia*, BC) supernatant (blue). We used either low iron availability (CAA + 100 µg/ml transferrin) (a) or high iron availability (CAA + 20uM $FeCl_3$) (b). PA was grown in 1 liter Erlenmeyer flasks in 200 ml medium at 37°C and 220 rpm. We followed growth by measuring the OD600 with a spectrophotometer (Ultrospec 2100 pro, Amersham Biosciences) until the cultures reached the mid- to late exponential phase, when we harvested the bacteria for following RNA isolation. We grew PA in 3 biological replicates per treatment. Each replicate of the PA and BC supernatant treatment contained a supernatant from an individually grown PA or BC monoculture under the respective iron condition. A fourth replicate was not harvested but was further monitored until the increase of OD600 levelled out, to control for the correct time point of harvesting.

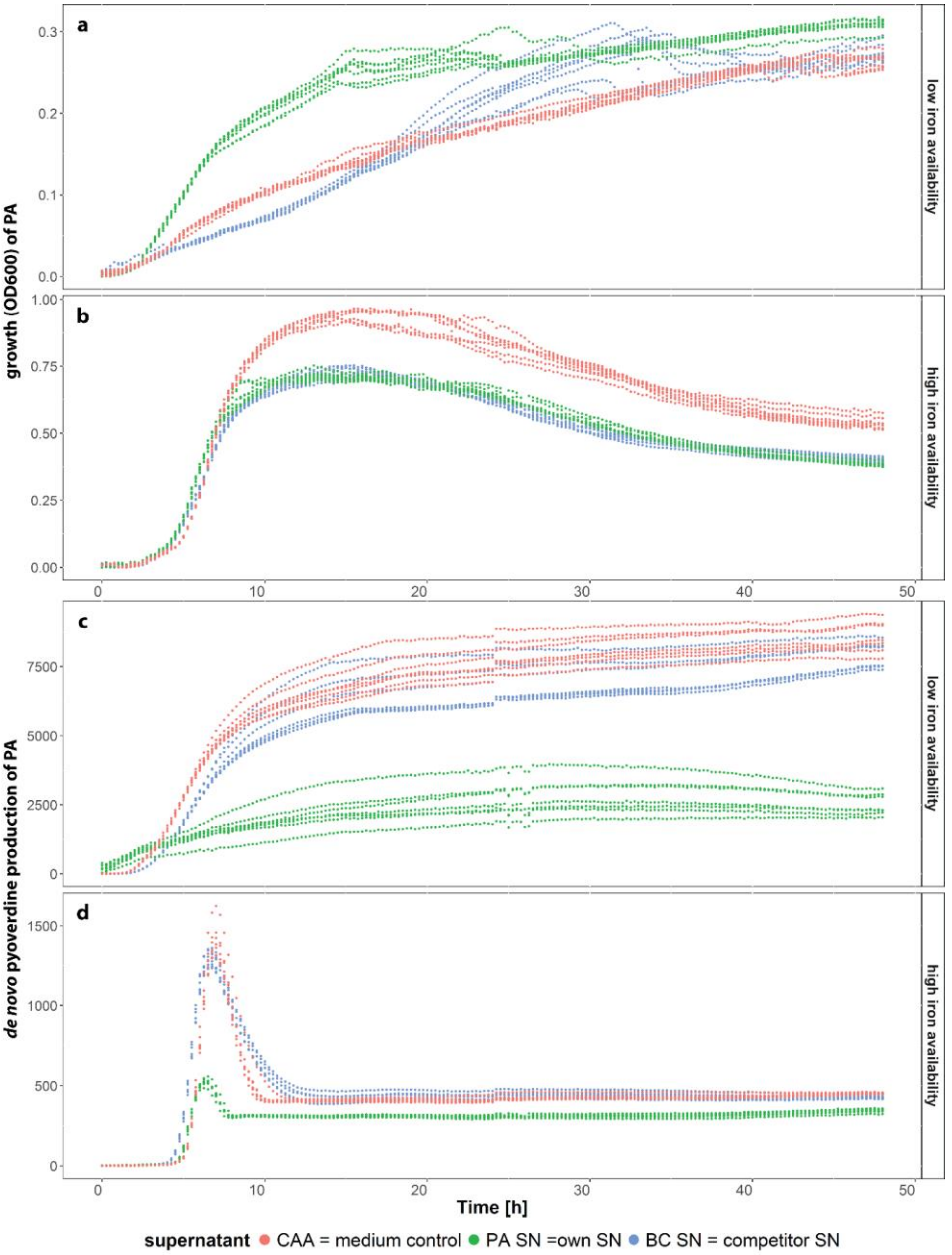


**Supplementary Fig. 2**: Growth and de novo pyoverdine production of *P. aeruginosa* (PA) grown in 100% fresh CAA medium (medium control, red), in 70% fresh CAA medium mixed with 30% of its own (PA) supernatant (green) or mixed with 30% of competitor (*B. cenocepacia*, BC) supernatant (blue) at low iron availability (a) and high iron availability (b). Under iron limitation (a), PA growth is stimulated by its own supernatant due to recyclable pyoverdine present in the supernatant. In contrast, the BC supernatant and the iron-limited CAA medium do not contain such growth stimulating siderophores. The BC supernatant slightly reduces PA growth compared to the other two treatments during early growth phases. Under iron-rich conditions (b), PA and BC supernatants mainly contain spent medium and no beneficial compounds, which reduces PA's overall growth compared to the medium control. The recyclable pyoverdine from the PA supernatant leads to a lower de novo pyoverdine synthesis by PA under iron limitation compared to the other two supernatant treatments (c). Under high iron availability, pyoverdine synthesis is low overall (d). The spent culture supernatant from PA suppresses PA's de novo pyoverdine synthesis to very low levels. Bacteria were cultured under two different iron regimes (low iron availability = CAA + 100 µg/ml transferrin; high iron availability = CAA + 20uM FeCl3). The OD600 and the autofluorescence of pyoverdine (excitation = 400 nm, emission = 460 nm) were measured over 48 hours of 7 replicates per treatment. The slight shifts in the pyoverdine fluorescence curves after 24h represent a technical artifact. This is because the plate reader protocol allowed only to program 24-hour runs, such that the protocol had to be manually relaunched after 24 hours, which caused the shift in fluorescence readings.

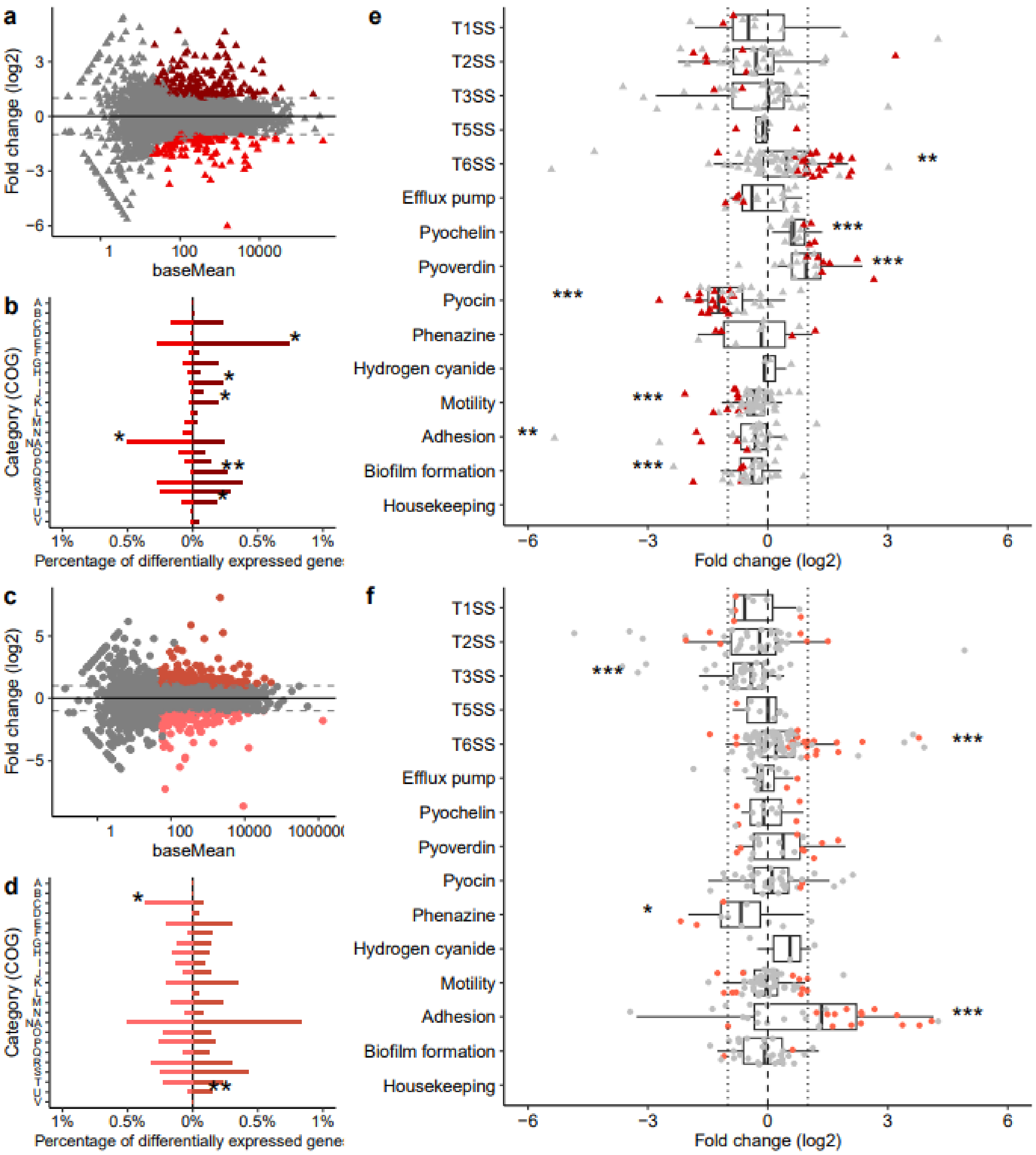


**Supplementary Fig. 3**: **Differential gene expression analysis of *P. aeruginosa* in response to its own spent supernatant.** The analysis compares the gene expression of *P. aeruginosa* growing in 70% of CAA medium + 30% of its own spent supernatant (SN.Pa) with the gene expression of *P. aeruginosa* growing in fresh CAA medium (SN.No), separately for iron-rich (intense shadings and triangles – a, b, e) and iron-limited (faint shadings and circles – c, d, f) conditions.

(a + c) MA-plots depicting significantly differentially regulated genes (in color) as a function of basemean values (FDR-$p_{adj}$ < 0.05 and log2-fold change > |1|).

(b + d) Percentage of differentially regulated genes according to their function (COG-categories). Asterisks indicate functional classes that are significantly over- or underrepresented based on the Fisher's Exact Test for count data comparing the number of all the expressed genes with the number of the significantly up- or downregulated genes. * $p < 0.05$, ** $p < 0.01$, *** $p < 0.001$. Capital letters stand for the following COG-categories; A: RNA processing and modification ; B: Chromatin structure and dynamics ; C: Energy production and conversion ; D: Cell cycle control, cell division, chromosome partitioning ; E: Amino acid transport and metabolism ; F: Nucleotide transport and metabolism ; G: Carbohydrate transport and metabolism ; H: Coenzyme transport and metabolism ; I: Lipid transport and metabolism ; J: Translation, ribosomal structure and biogenesis ; K: Transcription ; L: Replication, recombination and repair ; M: Cell wall/membrane/envelope biogenesis ; N: Cell motility ; NA: Unassigned ; O: Posttranslational modification, protein turnover, chaperones ; P: Inorganic ion transport and metabolism ; Q: Secondary metabolites biosynthesis, transport and catabolism ; R: General function prediction only ; S: Function unknown ; T: Signal transduction mechanisms ; U: Intracellular trafficking, secretion, and vesicular transport ; V: Defense mechanisms ; W: Extracellular structures ; X: Mobilome: prophages, transposons ; Y: Nuclear structure ; Z: Cytoskeleton.

(e + f) Data show the log2-fold change in gene expressions for gene clusters encoding competitive traits. Each dot represents an individual gene within a cluster. Colored and grey dots depict genes that are significantly and not-significantly differently regulated at the single gene level, respectively (FDR-adjusted p-values < 0.05). Asterisks indicate gene clusters that are, as a group, significantly up- and downregulated, according to a t-test conducted using zero as the expected mean (* $p < 0.05$, ** $p < 0.01$, *** $p < 0.001$). The dotted vertical lines represent log2-fold change = |1|.

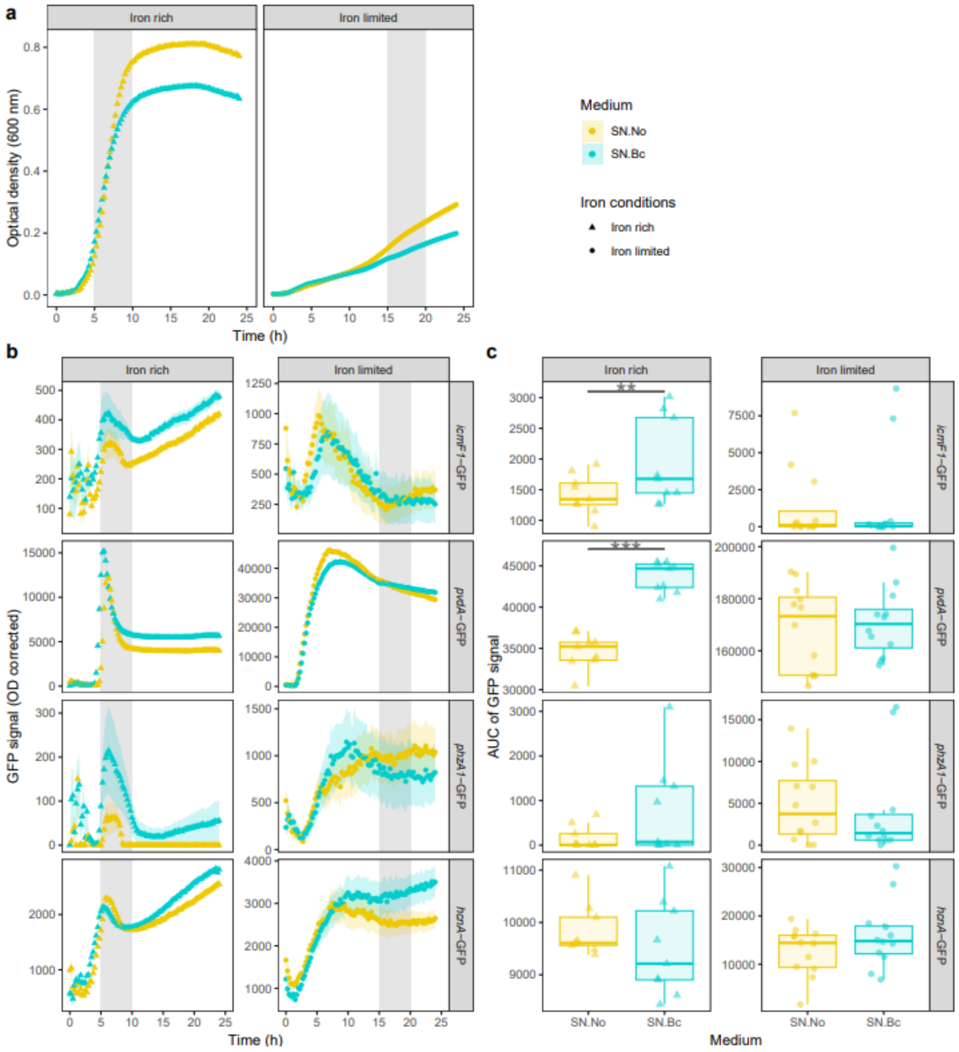


**Supplementary Fig. 4: Validation experiment #1, following the expression of four genes encoding competitive traits in *P. aeruginosa* under four experimental conditions using fluorescent gene-expression reporter strains.** *P. aeruginosa* bacteria were exposed to either fresh CAA medium (SN.No, yellow) or *B. cenocepacia* supernatant (SN.Bc, cyan, 30% spent supernatant + 70% CAA). Cultures were grown for 24 hours under both iron-rich (triangles) and iron-limited (circles) conditions. We used four reporter strains to track expression of genes encoding competitive traits (*icmF1* for structural component of the T6SS, *pvdA* for pyoverdine synthesis, *phzA1* for phenazine synthesis, and *hcnA* for hydrogen cyanide synthesis). We followed growth and gene expression of all strains over time in 9-12 independent replicates. Grey shaded areas highlight the time windows during which we harvested the cells for RNA extraction for the RNA-Seq experiment: between 5 and 10 hours in iron-rich medium, and between 15 and 20 hours in iron-limited medium.

(a) Growth curves of *P. aeruginosa* under the four different culturing conditions. The data depict the mean and standard error across all four reporter strains (with 9-12 replicates per strain). Standard errors are small and typically lay within the boundaries of the symbols. We observed that *P. aeruginosa* grows better in iron-rich compared to iron-limited conditions. In iron-rich conditions, all strains grew similarly until late exponential phase regardless of whether supernatant was supplemented or not. Towards the end of the exponential phase, *P.*

*aeruginosa* grew to higher yield in SN.No medium because more nutrients are present compared to the SN.Bc medium. In iron-limited conditions, the growth curves of *P. aeruginosa* start to diverge between SN.No and SN.Bc at intermediate time points of the growth phase.

(b) Temporal gene expression trajectories of the four selected genes in the four different culturing conditions. The data depict the mean and standard error calculated across all independent replicates. All GFP values are normalized by the corresponding OD values to account for growth differences across culturing conditions. In iron-rich conditions, we observed a peak of gene expression between the $5^{th}$ and $10^{th}$ hour (exponential growth phase). In iron-limited conditions, gene expression rose and then stayed stable or declined during the time window of interest (15-20 hours, shaded area).

(c) Statistical comparisons of the gene expression trajectories of the four selected genes. In order to compare the GFP reporter data with the RNA-Seq transcriptomic data, we calculated the area under the curve (AUC) of the GFP signal for the time period during which samples were collected for the RNA-Seq experiment (iron-rich conditions: 5-10 hours; iron-limited conditions: 15-20 hours). Please note that a perfect comparison of time points between the GFP-reporter experiment and the RNA-Seq experiment is difficult because the culturing conditions differed (reporter experiment: 96-well plates; RNA-Seq: 1 liter glass flasks) and so statistical analysis must be interpreted with caution. Individual data points depict independent replicates, and ANOVAs were used to compare the GFP signal levels between the two culturing conditions. ** and *** respectively for $p< 0.01$ and $p < 0.001$.

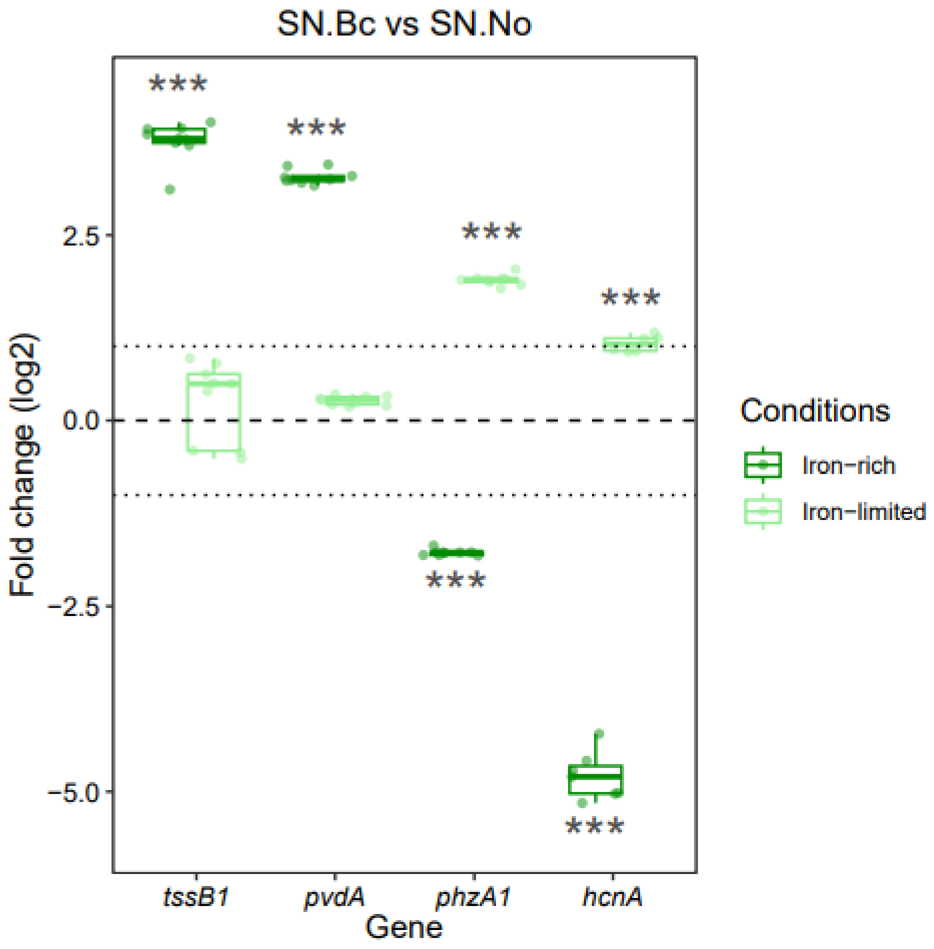


**Supplementary Fig. 5: Validation experiment #2 quantifying the differential expression of four genes encoding competitive traits in *P. aeruginosa* by qPCR.**

The expression of four genes (*tssB1*, for structural component of the T6SS, *pvdA* for pyoverdine synthesis, *phzA1* for phenazine synthesis, and *hcnA* for hydrogen cyanide synthesis) was measured by qPCR. Data show the log2-fold change in the expression of *P. aeruginosa* exposed to *B. cenocepacia* supernatant (SN.Bc) compared to fresh CAA medium (SN.No) in iron-rich (dark green) and iron-limited (light green) growth conditions. Each dot represents an individual replicate. Asterisks indicate genes that are significantly up- and downregulated according to a t-test conducted using zero as the expected mean (***p-value < 0.001). A mean log2-fold change higher than |1| was used as a threshold for biological relevance. These results confirm our main observations from Fig. 4. In response to *B. cenocepacia* supernatant exposure, *P. aeruginosa* upregulates genes encoding T6SS and pyoverdine production under iron-rich conditions, while it upregulates genes encoding phenazines and hydrogen cyanide synthesis under iron-limited conditions.

**Supplementary Table 1:** Number of reads mapped to the PAO1 reference genome per replicate.

| Nutrient condition | Supernatent treatment | Replicate | No. of reads |
|---|---|---|---|
| Iron limited | SN.No | 1 | 7924615 |
| | | 2 | 8592846 |
| | | 3 | 4988364 |
| | SN.Bc | 1 | 8827668 |
| | | 2 | 8049557 |
| | | 3 | 5954776 |
| | SN.Pa | 1 | 8183758 |
| | | 2 | 4975289 |
| | | 3 | 7651807 |
| Iron rich | SN.No | 1 | 8238867 |
| | | 2 | 6499571 |
| | | 3 | 5316425 |
| | SN.Bc | 1 | 6167446 |
| | | 2 | 6322462 |
| | | 3 | 6287578 |
| | SN.Pa | 1 | 6365862 |
| | | 2 | 4595376 |
| | | 3 | 6856674 |

**Supplementary Table 2:** Number of mapped and differentially expressed genes per comparison and growth condition in (a) absolute values, and (b) percentages.

While 5570 open reading frames were originally annotated in the *P. aeruginosa* PAO1 genome [1], this number has gone up to 5713 according to the *Pseudomonas* Genome Database (pseudomonas.com – DB Version 21.1 2022-11-20) [2] and the COG functional mapping database (2014 analysis) [3].

| Nutrient conditions | Comparison | Mapped genes | Differentially expressed genes (relaxed*) | Differentially expressed genes (stringent**) | Significantly down-regulated genes (stringent**) | Significantly up-regulated genes (stringent**) |
|---|---|---|---|---|---|---|
| **(a)** Total number of genes | | | | | | |
| Iron rich | SN.Bc vs SN.Pa | 5605 | 768 | 335 | 162 | 173 |
| | SN.Bc vs SN.No | 5606 | 1082 | 602 | 250 | 352 |
| | SN.Pa vs SN.No | 5610 | 617 | 319 | 126 | 193 |
| Iron limited | SN.Bc vs SN.Pa | 5626 | 559 | 327 | 158 | 169 |
| | SN.Bc vs SN.No | 5626 | 918 | 547 | 208 | 339 |
| | SN.Pa vs SN.No | 5624 | 844 | 439 | 195 | 244 |
| **(b)** Percentage of mapped genes | | | | | | |
| Iron rich | SN.Bc vs SN.Pa | 98.1 | 13.7 | 6.0 | 2.9 | 3.1 |
| | SN.Bc vs SN.No | 98.1 | 19.3 | 10.7 | 4.5 | 6.3 |
| | SN.Pa vs SN.No | 98.2 | 11.0 | 5.7 | 2.2 | 3.4 |
| Iron limited | SN.Bc vs SN.Pa | 98.5 | 9.9 | 5.8 | 2.8 | 3.0 |
| | SN.Bc vs SN.No | 98.5 | 16.3 | 9.7 | 3.7 | 6.0 |
| | SN.Pa vs SN.No | 98.4 | 15.0 | 7.8 | 3.5 | 4.3 |

* $p_{adj} < 0.05$

** $p_{adj} < 0.05$ and log2-fold > |1|

**Supplementary Table 3:** 50 top up- and down-regulated genes among the significantly differentially expressed genes by PA when grown in 70% CAA medium + 30% of BC supernatant (SN.Bc) compared to PA when grown in 100% CAA (SN.No), in iron-rich conditions.

| Locus tag | Gene name | Gene product | Category | log2-fold change | FDR-padj | |
|---|---|---|---|---|---|---|
| PA0883 | PA0883 | probable acyl-CoA lyase beta chain | G | 6.779417454 | 0.000749869 | top up-regulated genes |
| PA0880 | PA0880 | probable ring-cleaving dioxygenase | E | 6.716153888 | 5.6751E-123 | |
| PA0885 | dctQ | probable C4-dicarboxylate transporter | G | 6.248385641 | 9.45638E-70 | |
| PA0881 | PA0881 | hypothetical protein | R | 6.091987498 | 4.49347E-82 | |
| PA3501 | PA3501 | hypothetical protein | NA | 5.686782808 | 5.12408E-05 | |
| PA0884 | dctP | probable C4-dicarboxylate-binding periplasmic protein | G | 5.23482962 | 2.59175E-64 | |
| PA0882 | PA0882 | hypothetical protein | C | 5.223035388 | 1.02854E-58 | |
| PA0879 | PA0879 | probable acyl-CoA dehydrogenase | I | 4.727460426 | 1.36609E-90 | |
| **PA2386** | **pvdA** | **L-ornithine N5-oxygenase** | **Q** | **4.478017996** | **2.14395E-13** | |
| PA0886 | dctM | probable C4-dicarboxylate transporter | G | 4.157439035 | 1.39159E-39 | |
| PA2444 | glyA2 | serine hydroxymethyltransferase | E | 3.947806677 | 2.11363E-64 | |
| PA1258 | lhpM | Permease of ABC transporter, LhpM | E | 3.905332991 | 1.87358E-06 | |
| PA2385 | pvdQ | 3-oxo-C12-homoserine lactone acylase PvdQ | R | 3.81801979 | 3.56656E-11 | |
| PA2425 | pvdG | PvdG | Q | 3.757371838 | 0.024825732 | |
| PA0110 | PA0110 | hypothetical protein | S | 3.364543717 | 2.05056E-10 | |
| PA3502 | PA3502 | hypothetical protein | NA | 3.308307306 | 6.01662E-18 | |
| **PA0087** | **tssE1** | **TssE1** | **S** | **3.206482952** | **2.1642E-09** | |
| PA3441 | ssuF | probable molybdopterin-binding protein | H | 3.145449397 | 2.33792E-21 | |
| **PA0083** | **tssB1** | **TssB1** | **S** | **3.119318233** | **2.10419E-75** | |
| PA2452 | PA2452 | hypothetical protein | NA | 3.090461293 | 1.44016E-17 | |
| **PA2424** | **pvdL** | **PvdL** | **I / Q** | **3.086793568** | **4.53511E-29** | |
| **PA0089** | **tssG1** | **TssG1** | **S** | **3.037882069** | **2.46979E-25** | |
| PA2767 | PA2767 | probable enoyl-CoA hydratase/isomerase | I | 3.011449624 | 0.011030384 | |
| PA4501 | opdD | Glycine-glutamate dipeptide porin OpdP | NA | 2.989486727 | 6.91574E-24 | |
| PA2446 | gcvH2 | glycine cleavage system protein H2 | E | 2.905961692 | 5.80088E-31 | |
| **PA2402** | **pvdI** | **pyoverdine peptide synthetase** | **Q** | **2.888500039** | **2.17342E-29** | |
| PA0111 | PA0111 | hypothetical protein | NA | 2.878403095 | 6.17267E-12 | |
| PA1920 | nrdD | class III (anaerobic) ribonucleoside-triphosphate reductase subunit, NrdD | F / K | 2.872981141 | 2.42092E-40 | |
| PA1260 | lhpP | ABC transporter periplasmic-binding protein, LhpP | E / T | 2.804136561 | 2.39426E-08 | |
| PA2445 | gcvP2 | glycine cleavage system protein P2 | E | 2.784314806 | 1.82535E-43 | |
| PA4169 | PA4169 | conserved hypothetical protein | K | -6.648614947 | 0.000324779 | top down-regulated genes |
| PA2274 | PA2274 | hypothetical protein | NA | -6.518324677 | 0.00014415 | |
| PA5333 | PA5333 | conserved hypothetical protein | S | -5.406996641 | 0.002879912 | |
| PA1854 | PA1854 | conserved hypothetical protein | T | -5.052261217 | 0.011940957 | |
| PA1364 | PA1364 | probable transmembrane sensor | P / T | -4.879880471 | 0.015349115 | |
| PA1960 | PA1960 | hypothetical protein | S / K | -4.662838809 | 0.00501155 | |
| **PA1899** | **phzA2** | **probable phenazine biosynthesis protein** | **R** | **-4.361773211** | **7.04332E-08** | |
| PA4205 | mexG | hypothetical protein | S | -4.05614998 | 1.02862E-14 | |
| PA1906 | PA1906 | hypothetical protein | G / R / F | -3.85903577 | 2.05564E-11 | |
| PA2201 | PA2201 | hypothetical protein | R | -3.841310678 | 0.019845526 | |
| PA2300 | chiC | chitinase | G | -3.544354546 | 9.82861E-81 | |
| PA0512 | nirH | NirH | K | -3.473297856 | 2.21942E-06 | |
| **PA1900** | **phzB2** | **probable phenazine biosynthesis protein** | **R** | **-3.413983403** | **4.89408E-54** | |
| PA1907 | PA1907 | hypothetical protein | R | -3.34408065 | 5.24824E-12 | |
| **PA1901** | **phzC2** | **phenazine biosynthesis protein PhzC** | **E** | **-3.32868645** | **0.00149469** | |
| **PA4206** | **mexH** | **probable Resistance-Nodulation-Cell Division (RND) efflux membrane fusion protein precursor** | **M** | **-3.253176978** | **3.65613E-39** | |
| **PA3479** | **rhlA** | **rhamnosyltransferase chain A** | **I** | **-3.219323573** | **2.51277E-31** | |
| PA4377 | PA4377 | hypothetical protein | NA | -2.883341073 | 2.95879E-05 | |
| PA2064 | pcoB | copper resistance protein B precursor | P | -2.859719133 | 0.016209798 | |
| PA0808 | PA0808 | hypothetical protein | S | -2.818663767 | 0.027207757 | |

Putative traits involved in competition are written **in bold**.

Categories A: RNA processing and modification; B: Chromatin structure and dynamics; C: Energy production and conversion; D: Cell cycle control, cell division, chromosome partitioning; E: Amino acid transport and metabolism; F: Nucleotide transport and metabolism; G: Carbohydrate transport and metabolism; H: Coenzyme transport and metabolism; I: Lipid transport and metabolism; J: Translation, ribosomal structure and biogenesis; K: Transcription; L: Replication, recombination and repair; M: Cell wall/membrane/envelope biogenesis; N: Cell motility; NA: Non-assigned; O: Posttranslational modification, protein turnover, chaperones; P: Inorganic ion transport and metabolism; Q: Secondary metabolites biosynthesis, transport and catabolism; R: General function prediction only; S: Function unknown; T: Signal transduction mechanisms; U: Intracellular trafficking, secretion, and vesicular transport; V: Defense mechanisms; W: Extracellular structures; X: Mobilome (prophages, transposons); Y: Nuclear structure; Z: Cytoskeleton.

**Supplementary Table 4:** 50 top up- and down-regulated genes among the significantly differentially expressed genes by PA when grown in 70% CAA medium + 30% of BC supernatant (SN.Bc) compared to PA when grown in 100% CAA (SN.No), in iron-limited conditions.

| Locus tag | Gene name | Gene product | Category | log2-fold change | FDR-padj | |
|---|---|---|---|---|---|---|
| PA0574.1 | PA0574.1 | tRNA-Met | NA | 5.992365211 | 0.002630217 | **top up-regulated genes** |
| PA5084 | dguA | DguA | E | 5.548005282 | 3.22137E-11 | |
| PA2031 | PA2031 | hypothetical protein | NA | 5.272347354 | 2.55553E-82 | |
| PA5083 | dguB | Rid2 subfamily protein | J | 5.259484405 | 1.74204E-42 | |
| PA5351 | rubA1 | Rubredoxin 1 | C | 5.233073313 | 0.031255851 | |
| PA2030 | PA2030 | hypothetical protein | NA | 4.98593747 | 9.15726E-61 | |
| PA5081 | PA5081 | hypothetical protein | F | 4.696293662 | 2.56062E-09 | |
| PA5082 | dguC | DguC | E / T | 4.613201224 | 7.1094E-09 | |
| PA2634 | aceA | isocitrate lyase AceA | C | 4.533768697 | 6.1869E-119 | |
| PA0582 | folB | dihydroneopterin aldolase | H | 4.349851713 | 0.023508487 | |
| PA1983 | exaB | cytochrome c550 | C | 4.320346983 | 0.018536164 | |
| PA3779 | PA3779 | putative periplasmic substrate binding protein | G | 4.032200374 | 1.27525E-19 | |
| PA2225 | PA2225 | hypothetical protein | NA | 3.997729717 | 0.014092963 | |
| PA2228 | PA2228 | hypothetical protein | V | 3.987402845 | 0.005042686 | |
| **PA4650** | **cupE3** | **Pilin subunit CupE3** | **S** | **3.900932494** | **4.43325E-21** | |
| **PA4649** | **cupE2** | **Pilin subunit CupE2** | **S** | **3.793262433** | **5.41175E-17** | |
| PA0915 | yehS | conserved hypothetical protein | S | 3.582104349 | 0.029022588 | |
| PA3233 | PA3233 | hypothetical protein | T | 3.326489568 | 1.39585E-29 | |
| PA5550 | glmR | GlmR transcriptional regulator | G / K | 3.23273089 | 2.44309E-11 | |
| **PA4651** | **cupE4** | **Pilin assembly chaperone CupE4** | **N / U** | **3.222785236** | **1.31849E-13** | |
| PA1313 | PA1313 | probable major facilitator superfamily (MFS) transporter | G | 3.211145348 | 0.016108968 | |
| **PA4648** | **cupE1** | **Pilin subunit CupE1** | **S** | **3.188403815** | **1.34966E-33** | |
| PA5445 | psecoA | probable coenzyme A transferase | C | 3.170832043 | 3.33547E-40 | |
| PA2027 | PA2027 | hypothetical protein | NA | 2.963210764 | 0.011048609 | |
| PA3234 | yjcG | probable sodium:solute symporter | R | 2.895763937 | 1.21295E-25 | |
| PA3235 | yjcH | conserved hypothetical protein | S | 2.88523229 | 0.000831955 | |
| PA2029 | PA2029 | hypothetical protein | S | 2.817318639 | 3.23956E-12 | |
| PA4857 | tspR | TspR | U | 2.784230003 | 0.011691392 | |
| PA3305.1 | phrS | PhrS | NA | -6.304343719 | 1.01855E-09 | **top down-regulated genes** |
| PA0700 | PA0700 | hypothetical protein | NA | -5.954225031 | 0.009509578 | |
| PA3750 | PA3750 | hypothetical protein | E | -5.025520808 | 0.016649845 | |
| PA2759 | PA2759 | hypothetical protein | NA | -4.991949127 | 2.09632E-34 | |
| PA4359 | feoA | conserved hypothetical protein | P | -4.627188264 | 0.007439029 | |
| PA3431 | ywbG | conserved hypothetical protein | M | -4.594945605 | 1.49616E-22 | |
| PA5027 | PA5027 | hypothetical protein | T | -4.472664867 | 0.000191562 | |
| PA0200 | PA0200 | hypothetical protein | NA | -4.33657223 | 2.25259E-12 | |
| PA3006 | psrA | transcriptional regulator PsrA | K | -4.334296887 | 6.15714E-06 | |
| PA3432 | PA3432 | hypothetical protein | R | -4.309115914 | 2.26721E-20 | |
| PA3502 | PA3502 | hypothetical protein | NA | -4.173967503 | 0.015359044 | |
| PA1673 | PA1673 | hypothetical protein | P | -3.707528347 | 2.91264E-07 | |
| PA5475 | PA5475 | hypothetical protein | R / K | -3.593819664 | 4.57763E-12 | |
| PA4067 | oprG | | M | -3.350659629 | 2.16559E-06 | |
| PA2753 | PA2753 | hypothetical protein | NA | -3.308456523 | 4.20745E-10 | |
| PA4352 | PA4352 | conserved hypothetical protein | T | -3.176892498 | 3.36124E-11 | |
| PA2461 | PA2461 | hypothetical protein | S | -3.091404021 | 0.002775268 | |
| PA1395 | PA1395 | hypothetical protein | NA | -3.04297153 | 0.017273502 | |
| PA0273 | PA0273 | probable major facilitator superfamily (MFS) transporter | P | -2.990261096 | 0.002143502 | |
| PA1546 | hemN | oxygen-independent coproporphyrinogen III oxidase | H | -2.786747695 | 1.42578E-13 | |
| PA4917 | nadD2 | nicotinate mononucleotide adenylyltransferase NadD2 | H | -2.675846742 | 0.0000939 | |
| PA4350 | olsB | OlsB | R | -2.670090289 | 0.014720423 | |

Putative traits involved in competition are written **in bold**.

Categories A: RNA processing and modification; B: Chromatin structure and dynamics; C: Energy production and conversion; D: Cell cycle control, cell division, chromosome partitioning; E: Amino acid transport and metabolism; F: Nucleotide transport and metabolism; G: Carbohydrate transport and metabolism; H: Coenzyme transport and metabolism; I: Lipid transport and metabolism; J: Translation, ribosomal structure and biogenesis; K: Transcription; L: Replication, recombination and repair; M: Cell wall/membrane/envelope biogenesis; N: Cell motility; NA: Non-assigned; O: Posttranslational modification, protein turnover, chaperones; P: Inorganic ion transport and metabolism; Q: Secondary metabolites biosynthesis, transport and catabolism; R: General function prediction only; S: Function unknown; T: Signal transduction mechanisms; U: Intracellular trafficking, secretion, and vesicular transport; V: Defense mechanisms; W: Extracellular structures; X: Mobilome (prophages, transposons); Y: Nuclear structure; Z: Cytoskeleton.

**Supplementary Table 5:** 50 top up- and down-regulated genes among the significantly differentially expressed genes by PA when grown in 70% CAA medium + 30% of BC supernatant (SN.Bc) compared to PA when grown in 70% CAA + 30% of its own supernatant (SN.Pa), in iron-rich conditions.

| Locus tag | Gene name | Gene product | Category | log2-fold change | FDR-padj | |
|---|---|---|---|---|---|---|
| PA1418 | PA1418 | probable sodium:solute symport protein | E / R | 5.001626292 | 0.000201627 | top up-regulated genes |
| PA1420 | PA1420 | hypothetical protein | S | 3.829425706 | 0.003262037 | |
| PA1417 | PA1417 | probable decarboxylase | E / H | 3.787517064 | 0.001510054 | |
| PA3441 | ssuF | probable molybdopterin-binding protein | H | 3.473759577 | 7.67316E-21 | |
| PA0885 | dctQ | probable C4-dicarboxylate transporter | G | 3.192268679 | 2.00691E-30 | |
| PA1416 | PA1416 | conserved hypothetical protein | C | 3.185038065 | 0.000474132 | |
| PA0194 | PA0194 | hypothetical protein | Q | 3.030452489 | 0.000017824 | |
| PA3235 | yjcH | conserved hypothetical protein | S | 3.01490138 | 0.000015561 | |
| PA3038 | opdQ | OpdQ | NA | 2.979973003 | 2.62675E-29 | |
| PA3864 | dauR | Transcriptional regulator of the dauBAR operon, DauR | S | 2.80536383 | 2.61531E-05 | |
| PA0881 | PA0881 | hypothetical protein | R | 2.792690266 | 7.62206E-28 | |
| PA0883 | PA0883 | probable acyl-CoA lyase beta chain | G | 2.773819982 | 6.15165E-21 | |
| PA0880 | PA0880 | probable ring-cleaving dioxygenase | E | 2.691780831 | 8.43005E-47 | |
| PA3865 | PA3865 | putative periplasmic lysine-, arginine-, ornithine-binding protein | E / T | 2.632041459 | 9.50054E-14 | |
| **PA0637** | **PA0637** | **conserved hypothetical protein** | **S** | **2.563680939** | **0.000189379** | |
| PA0882 | PA0882 | hypothetical protein | C | 2.5015987 | 4.52143E-20 | |
| PA3234 | yjcG | probable sodium:solute symporter | R | 2.484993553 | 9.08718E-05 | |
| PA1421 | gbuA | guanidinobutyrase | E | 2.483437576 | 0.032053011 | |
| PA0884 | dctP | probable C4-dicarboxylate-binding periplasmic protein | G | 2.47256403 | 1.38756E-16 | |
| PA0886 | dctM | probable C4-dicarboxylate transporter | G | 2.452144651 | 9.28192E-16 | |
| PA3863 | dauA | FAD-dependent catabolic D-arginine dehydrogenase, DauA | E | 2.409129985 | 7.23525E-08 | |
| PA3233 | PA3233 | hypothetical protein | T | 2.40504142 | 4.97868E-13 | |
| PA0879 | PA0879 | probable acyl-CoA dehydrogenase | I | 2.401448197 | 1.13776E-29 | |
| PA3445 | PA3445 | conserved hypothetical protein | P | 2.353429146 | 2.2948E-12 | |
| PA3502 | PA3502 | hypothetical protein | NA | 2.272764762 | 1.47334E-07 | |
| PA3232 | PA3232 | probable nuclease | L | 2.244893311 | 2.04544E-07 | |
| PA2313 | PA2313 | hypothetical protein | NA | 2.22895406 | 2.79302E-06 | |
| PA2085 | PA2085 | probable ring-hydroxylating dioxygenase small subunit | Q | 2.094597391 | 4.34371E-05 | |
| PA0098 | PA0098 | hypothetical protein | I / Q | 2.061049295 | 2.08959E-06 | |
| PA2393 | PA2393 | putative dipeptidase | E | 2.047201153 | 0.036903809 | |
| PA1920 | nrdD | class III (anaerobic) ribonucleoside-triphosphate reductase subunit, NrdD | F / K | 2.030955613 | 2.01191E-16 | |
| PA4149 | acoX | conserved hypothetical protein | S | -3.459526617 | 4.20224E-07 | top down-regulated genes |
| PA1906 | PA1906 | hypothetical protein | G / R / F | -3.162937277 | 0.000206424 | |
| PA4150 | acoA | probable dehydrogenase E1 component | C | -3.114848007 | 0.014872185 | |
| PA4205 | mexG | hypothetical protein | S | -3.0672209 | 4.95927E-05 | |
| PA5180 | fdhD | conserved hypothetical protein | C | -2.925436273 | 2.05016E-14 | |
| PA4364 | PA4364 | hypothetical protein | S | -2.894162308 | 2.53985E-05 | |
| PA3721 | nalC | NalC | K | -2.880618138 | 0.001021299 | |
| PA1325 | yybH | conserved hypothetical protein | S | -2.866357507 | 0.002432697 | |
| PA3888 | opuCD | OpuC ABC transporter, permease protein, OpuCD | E | -2.634644126 | 0.004004641 | |
| PA5181 | PA5181 | probable oxidoreductase | C | -2.581349399 | 4.70405E-22 | |
| PA4181 | PA4181 | hypothetical protein | S | -2.466083417 | 1.29131E-14 | |
| **PA4206** | **mexH** | **probable Resistance-Nodulation-Cell Division (RND) efflux membrane fusion protein precursor** | **M** | **-2.30883302** | **1.32464E-21** | |
| **PA1900** | **phzB2** | **probable phenazine biosynthesis protein** | **R** | **-2.235881952** | **2.88143E-11** | |
| PA4183 | PA4183 | hypothetical protein | R | -2.231548249 | 5.02965E-07 | |
| PA5507 | PA5507 | hypothetical protein | Q | -2.229077659 | 5.17147E-10 | |
| PA3189 | gltF | probable permease of ABC sugar transporter | G | -2.18030371 | 4.76135E-09 | |
| PA3720 | PA3720 | hypothetical protein | NA | -2.178785103 | 2.48567E-07 | |
| PA4365 | lysE | Lysine efflux permease | R | -2.105569285 | 9.86636E-07 | |
| PA3190 | gltB | probable binding protein component of ABC sugar transporter | G | -2.091737301 | 1.48915E-17 | |

Putative traits involved in competition are written **in bold**.

Categories A: RNA processing and modification; B: Chromatin structure and dynamics; C: Energy production and conversion; D: Cell cycle control, cell division, chromosome partitioning; E: Amino acid transport and metabolism; F: Nucleotide transport and metabolism; G: Carbohydrate transport and metabolism; H: Coenzyme transport and metabolism; I: Lipid transport and metabolism; J: Translation, ribosomal structure and biogenesis; K: Transcription; L: Replication, recombination and repair; M: Cell wall/membrane/envelope biogenesis; N: Cell motility; NA: Non-assigned; O: Posttranslational modification, protein turnover, chaperones; P: Inorganic ion transport and metabolism; Q: Secondary metabolites biosynthesis, transport and catabolism; R: General function prediction only; S: Function unknown; T: Signal transduction mechanisms; U: Intracellular trafficking, secretion, and vesicular transport; V: Defense mechanisms; W: Extracellular structures; X: Mobilome (prophages, transposons); Y: Nuclear structure; Z: Cytoskeleton.

**Supplementary Table 6:** 50 top up- and down-regulated genes among the significantly differentially expressed genes by PA when grown in 70% CAA medium + 30% of BC supernatant (SN.Bc) compared to PA when grown in 70% CAA + 30% of its own supernatant (SN.Pa), in iron-limited conditions.

| Locus tag | Gene name | Gene product | Category | log2-fold change | FDR-padj | |
|---|---|---|---|---|---|---|
| PA5084 | dguA | DguA | E | 6.085610142 | 1.68664E-17 | top up-regulated genes |
| PA5083 | dguB | Rid2 subfamily protein | J | 5.931882012 | 1.19077E-21 | |
| PA5082 | dguC | DguC | E / T | 5.345643457 | 2.27802E-14 | |
| PA2031 | PA2031 | hypothetical protein | NA | 4.838240676 | 1.1626E-111 | |
| PA5081 | PA5081 | hypothetical protein | F | 4.674834164 | 1.57906E-09 | |
| PA2030 | PA2030 | hypothetical protein | NA | 4.56086605 | 1.62222E-59 | |
| PA0755 | opdH | cis-aconitate porin OpdH | NA | 4.320397488 | 1.61197E-47 | |
| PA5180 | fdhD | conserved hypothetical protein | C | 3.500164966 | 2.93612E-39 | |
| PA2029 | PA2029 | hypothetical protein | S | 3.332088165 | 3.18196E-17 | |
| PA0195.1 | pntAB | putative NAD(P) transhydrogenase, subunit alpha part 2 | NA | 3.058501921 | 1.92027E-17 | |
| PA3235 | yjcH | conserved hypothetical protein | S | 2.998590091 | 1.79061E-15 | |
| PA3569 | mmsB | 3-hydroxyisobutyrate dehydrogenase | I | 2.948664718 | 2.52836E-20 | |
| PA2004 | PA2004 | conserved hypothetical protein | NA | 2.727881623 | 1.08013E-25 | |
| PA3570 | mmsA | methylmalonate-semialdehyde dehydrogenase | C | 2.715389222 | 4.88529E-26 | |
| PA3781 | PA3781 | probable transporter | G | 2.704985186 | 4.00247E-07 | |
| PA2003 | bdhA | 3-hydroxybutyrate dehydrogenase | R / I / Q | 2.667858462 | 9.49014E-29 | |
| PA5549 | glmS | glucosamine--fructose-6-phosphate aminotransferase | M | 2.644093707 | 3.69747E-11 | |
| PA3365 | amiB | probable chaperone | O | 2.622054201 | 2.95466E-18 | |
| PA0754 | PA0754 | hypothetical protein | S | 2.578136536 | 1.67114E-14 | |
| PA3234 | yjcG | probable sodium:solute symporter | R | 2.565418123 | 1.4932E-18 | |
| PA3233 | PA3233 | hypothetical protein | T | 2.56143509 | 1.15636E-18 | |
| PA5550 | glmR | GlmR transcriptional regulator | G / K | 2.555116777 | 5.79338E-09 | |
| PA3190 | gltB | probable binding protein component of ABC sugar transporter | G | 2.55039658 | 0.009106517 | |
| PA3779 | PA3779 | putative periplasmic substrate binding protein | G | 2.546527472 | 7.2804E-13 | |
| PA3038 | opdQ | OpdQ | NA | 2.523078812 | 1.90747E-36 | |
| PA2634 | aceA | isocitrate lyase AceA | C | 2.517131839 | 1.28795E-25 | |
| PA3710 | PA3710 | probable GMC-type oxidoreductase | E | 2.455605475 | 7.00403E-19 | |
| PA3189 | gltF | probable permease of ABC sugar transporter | G | 2.401458449 | 1.5692E-06 | |
| PA0865 | hpd | 4-hydroxyphenylpyruvate dioxygenase | E / R | 2.380087273 | 1.2942E-21 | |
| PA3305.1 | phrS | PhrS | NA | 2.319701092 | 4.92973E-05 | |
| PA0049 | PA0049 | hypothetical protein | NA | -7.481283312 | 1.07342E-11 | top down-regulated genes |
| PA2116 | PA2116 | conserved hypothetical protein | S | -5.857058701 | 0.002091496 | |
| PA2111 | PA2111 | hypothetical protein | E | -5.432078007 | 2.14525E-33 | |
| PA2110 | PA2110 | hypothetical protein | E | -5.310022996 | 0.024185742 | |
| PA2114 | PA2114 | probable major facilitator superfamily (MFS) transporter | E / G / R / P | -5.292242512 | 0.00039768 | |
| PA2113 | opdO | pyroglutamate porin OpdO | NA | -4.312663724 | 0.031279675 | |
| **PA5267** | **hcpB** | **secreted protein Hcp** | **S** | **-4.000156817** | **5.73952E-15** | |
| PA0048 | PA0048 | probable transcriptional regulator | K | -3.374428712 | 1.42548E-05 | |
| PA2112 | PA2112 | conserved hypothetical protein | R | -3.164066659 | 1.37801E-07 | |
| PA1337 | ansB | glutaminase-asparaginase | E / J | -2.887909581 | 3.15231E-38 | |
| PA2444 | glyA2 | serine hydroxymethyltransferase | E | -2.834739381 | 8.99513E-27 | |
| PA1342 | aatJ | putative acidic amino acid ABC transporter substrate-binding protein | E / T | -2.755875404 | 5.42949E-37 | |
| PA1654 | PA1654 | probable aminotransferase | E /K | -2.721092945 | 1.94642E-29 | |
| **PA4175** | **piv** | **protease IV** | **NA** | **-2.428471448** | **1.59476E-30** | |
| PA1341 | aatQ | AatQ | E | -2.424847299 | 1.38233E-11 | |
| PA1338 | ggt | gamma-glutamyltranspeptidase precursor | E | -2.413684442 | 7.47425E-25 | |
| PA3431 | ywbG | conserved hypothetical protein | M | -2.391862854 | 0.000882558 | |
| PA2446 | gcvH2 | glycine cleavage system protein H2 | E | -2.370799866 | 5.27664E-23 | |
| **PA1665** | **fha2** | **Fha2** | **T** | **-2.365949168** | **3.59496E-11** | |
| PA5507 | PA5507 | hypothetical protein | Q | -2.341806472 | 3.43958E-12 | |

Putative traits involved in competition are written **in bold**.

Categories A: RNA processing and modification; B: Chromatin structure and dynamics; C: Energy production and conversion; D: Cell cycle control, cell division, chromosome partitioning; E: Amino acid transport and metabolism; F: Nucleotide transport and metabolism; G: Carbohydrate transport and metabolism; H: Coenzyme transport and metabolism; I: Lipid transport and metabolism; J: Translation, ribosomal structure and biogenesis; K: Transcription; L: Replication, recombination and repair; M: Cell wall/membrane/envelope biogenesis; N: Cell motility; NA: Non-assigned; O: Posttranslational modification, protein turnover, chaperones; P: Inorganic ion transport and metabolism; Q: Secondary metabolites biosynthesis, transport and catabolism; R: General function prediction only; S: Function unknown; T: Signal transduction mechanisms; U: Intracellular trafficking, secretion, and vesicular transport; V: Defense mechanisms; W: Extracellular structures; X: Mobilome (prophages, transposons); Y: Nuclear structure; Z: Cytoskeleton.

**Supplementary Table 7- 10 are shown in separate Excel sheets.**

**Supplementary Table 7:** List of PA genes encoding competitive traits and regulatory systems. The list is not exhaustive but covers a broad spectrum of competitive traits including secretion systems, efflux pumps, siderophores, toxins, motility, adhesion and biofilm, and regulatory systems including elements of the general stress response, quorum sensing systems and two-component signalling systems.

**Supplementary Table 8:** Transcriptional analysis of PA genes encoding competitive traits and regulatory systems (see Supplementary table 7) in PA when grown in 70% CAA medium + 30% of BC supernatant (SN.Bc) compared to PA when grown in 100% CAA (SN.No), in iron-rich and iron-limited conditions. Significantly differentially expressed genes are highlighted in bold (FDR-$p_{adj}$ < 0.05).

**Supplementary Table 9:** Transcriptional analysis of PA genes encoding competitive traits and regulatory systems (see Supplementary table 7) in PA when grown in 70% CAA medium + 30% of BC supernatant (SN.Bc) compared to PA when grown in 70% CAA + 30% of its own supernatant (SN.Pa), in iron-rich and iron-limited conditions. Significantly differentially expressed genes are highlighted in bold (FDR-$p_{adj}$ < 0.05).

**Supplementary Table 10:** Transcriptional analysis of PA genes encoding competitive traits and regulatory systems (see Supplementary table 7) in PA when grown in 70% CAA medium + 30% of its own supernatant (SN.Pa) compared to PA when grown in 100% CAA (SN.No), in iron-rich and iron-limited conditions. Significantly differentially expressed genes are highlighted in bold (FDR-$p_{adj}$ < 0.05).

**Supplementary Table 11:** Statistical parameters resulting from t-tests performed on the competitive gene clusters. The mean of the log2-fold changes for each functional category was compared to the theoretical value of zero using one-sample t-test. The categories that are, as a group, significantly up- and down-regulated in *P. aeruginosa* upon exposure to *B. cenocepacia* supernatant are written in red and blue respectively.

**Supplementary References:**


1. Stover, C. K. *et al.* Complete genome sequence of *Pseudomonas aeruginosa* PAO1, an opportunistic pathogen. *Nature* **406**, 959–964 (2000).
2. Winsor, G. L. *et al.* Enhanced annotations and features for comparing thousands of *Pseudomonas* genomes in the Pseudomonas genome database. *Nucleic Acids Res.* **44**, D646-653 (2016).
3. Galperin, M. Y., Makarova, K. S., Wolf, Y. I. & Koonin, E. V. Expanded microbial genome coverage and improved protein family annotation in the COG database. *Nucleic Acids Res.* **43**, D261-269 (2015).
